# Preliminary Characterization of Phage-like Particles from the Male-Killing Mollicute *Spiroplasma poulsonii* (an Endosymbiont of *Drosophila*)

**DOI:** 10.1101/2021.12.09.471767

**Authors:** Paulino Ramirez, Justin C. Leavitt, Jason J. Gill, Mariana Mateos

**Author notes:** University of Texas Health Science Center-San Antonio, San Antonio, TX, USA. **Author contributions:** Conceived and designed the analysis: PR, JCL, JJG, MM Collected the data: PR, JCL, JJG Contributed materials, data or analysis tools: PR, JCL, JJG, MM Performed the analyses and interpreted the results: PR, JCL, JJG, MM, Wrote the paper: PR, MM Edited the paper: PR, MM, JCL, JJG. **Data Availability Statement** The raw sequence data used for the present study are available at NCBI under Project Number PRJNA545743; BioSamples SAMN11919470, SAMN11919470, SAMN23459927; and SRA SRR17050036, SRR17063333, SRR17065466. Assembled and annotated contigs are available at NCBI under GenBank Accession Nos. OL689226-OL689230 and OL778852.

## Abstract

Bacteriophages are vastly abundant, diverse, and influential, but with few exceptions (e.g. the Proteobacteria genera *Wolbachia* and *Hamiltonella*), the role of phages in heritable bacteria-arthropod interactions, which are ubiquitous and diverse, remains largely unexplored. Despite prior studies documenting phage-like particles in the mollicute *Spiroplasma* associated with *Drosophila* flies, genomic sequences of such phage are lacking, and their effects on the *Spiroplasma*-*Drosophila* interaction have not been comprehensively characterized. We used a density step gradient to isolate phage-like particles from the male-killing bacterium *Spiroplasma poulsonii* (strains NSRO and MSRO-Br) harbored by *Drosophila melanogaster*. Isolated particles were subjected to DNA sequencing, assembly, and annotation. Several lines of evidence suggest that we recovered phage-like particles of similar features (shape, size, DNA content) to those previously reported in *Drosophila*-associated *Spiroplasma* strains. We recovered three ∼19 kb phage-like contigs (two in NSRO and one in MSRO-Br) containing 21–24 open reading frames, a read-alignment pattern consistent with circular permutation, and terminal redundancy (at least in NSRO). Although our results do not allow us to distinguish whether these phage-like contigs represent infective phage-like particles capable of transmitting their DNA to new hosts, their encoding of several typical phage genes suggests that they are at least remnants of functional phage. We also recovered two smaller non-phage-like contigs encoding a known *Spiroplasma* toxin (Ribosome Inactivating Protein; RIP), and an insertion element, suggesting that they are packaged into particles. Substantial homology of our particle-derived contigs was found in the genome assemblies of members of the *Spiroplasma poulsonii* clade.

## Introduction

Viruses of bacteria, referred to as bacteriophages or phages, are the most abundant and diverse biological entities on the planet. They exert extraordinary impacts on ecosystems, by controlling bacterial abundance, shaping community composition, and altering biogeochemical cycling. Phages also mediate horizontal gene transfer and influence their host’s ability to invade and proliferate in eukaryotes [reviewed in 1]. Many phages carry toxin-encoding genes that are responsible for the virulence of their host bacteria [2-4], including those bacteria engaged in intimate heritable associations with arthropod hosts [5].

Heritable associations between bacteria and arthropods are ubiquitous, diverse, and influential [6], and can be classified under two categories. Obligate or primary associations are those in which the bacterium is absolutely required for host survival/reproduction, typically conferring a nutritional benefit, vertical transmission is perfect (i.e., females transmit the bacterium to 100% of their offspring), and infection is effectively fixed in the host population. On the other hand, facultative or secondary associations are those in which the bacterium is not required for host survival, vertical transmission is usually imperfect, and infection is variable in the host population. Facultative endosymbionts tend to manipulate host reproduction to their own benefit, or confer a conditional fitness advantage to hosts [e.g. tolerance to environmental stressors; 7] or defense against natural enemies [reviewed in 8].

Recent studies of the mechanistic bases of arthropod-bacteria reproductive manipulation and defensive mutualisms have implicated phages. For example, members of the Alphaproteobacteria genus *Wolbachia* (the most common facultative heritable symbiont of arthropods), cause several reproductive phenotypes including cytoplasmic incompatibility (CI). CI is a type of reproductive manipulation that leads to embryo lethality in the offspring of *Wolbachia-*infected males and uninfected females [9, 10]. The *Wolbachia*-encoded genes responsible for CI were recently discovered and are encoded by a temperate (= lysogenic) phage known as “WO” [11-14]. Similarly, the male-killing phenotype of certain *Wolbachia* strains also appears to be encoded by a phage [15]. Phages have also been implicated in the defense phenotype conferred by certain bacteria to their arthropod hosts. Strains of *Hamiltonella defensa* (Gammaproteobacteria) that are infected with a particular phage (APSE3) produce a soluble factor that prevents successful development of a parasitic wasp of the aphid [16]. Therefore, such phage-infected *H. defensa* strains are defensive mutualists of their aphid host.

Beyond *Wolbachia* and *Hamiltonella*, numerous other facultative heritable bacteria are commensals, reproductive parasites, and/or mutualists of arthropods [reviewed by 6]. One such common bacterial lineage is *Spiroplasma* (Mollicutes), a diverse genus infecting many eukaryotic hosts, particularly arthropods and plants, inhabiting both intra-(e.g. phloem, ovaries, salivary glands) and extra-cellular environments (e.g. hemolymph). *Spiroplasma*-host interactions range from parasitic to mutualistic [e.g. 17, 18, reviewed by 19], and transmission modes are either predominantly vertical (as in *Drosophila* flies) or horizontal [i.e., from the environment or via a vector; 20]. Four independent clades of *Spiroplasma* are associated with members of the genus *Drosophila* [21]. The best studied of these is the *Spiroplasma poulsonii* clade, which includes strains that associate naturally with several divergent species of *Drosophila*, including the model organism *D. melanogaster* (Table 1) [reviewed by 22, 23]. *Spiroplasma* is abundant in the hemolymph of its *Drosophila* host, from where it has been easily transferred to diverse recipient *Drosophila* species, often achieving vertical transmission in the new host [24]. Several strains of *S. poulsonii* are male killers (referred to as Sex Ratio Organisms, SROs or SRs; Table 1), whereby male *Drosophila* die during early embryonic development [25]. In addition, all strains of *S. poulsonii* examined to date prevent the successful development of at least one parasitoid wasp species of *Drosophila* [Table 1; reviewed in 26]. Collectively, numerous studies on *poulsonii*-clade strains have revealed that their genomes evolve extremely rapidly [27], and have uncovered *Drosophila* and *Spiroplasma* factors and mechanisms underlying vertical transmission, male killing, and defense against wasps and nematodes [28-40], including several *Spiroplasma*-encoded toxins, such as Ribosome Inactivating Proteins (RIPs). The possible role of active phages in *Drosophila*-*Spiroplasma* interactions, however, has not been addressed.

**Table 1.** *Drosophila*-associated *Spiroplasma* strains belonging to the *poulsonii* clade and/or in which phage-like particles have been reported.

| *poulsonii* clade | "Modern" Abbreviation<sup>b</sup> | native *Drosophila* host | Male-killer ("Old" abbreviation<sup>d</sup>) | defensive of *Drosophila*: natural enemy affected | In vitro culture | Genome sequence | Past studies: Particles reported, isolation method | Strains targeted in the present study and types of data obtained |
| --- | --- | --- | --- | --- | --- | --- | --- | --- |
| yes | sNeb | *D. nebulosa* | yes<sup>c</sup> (NSRO) | unknown | no | no | yes, first degree<sup>g</sup> | particles isolation, TEM, short- and long-read sequencing |
| yes | sWil | *D. willistoni* | yes<sup>c</sup> (WSRO) | unknown | yes<sup>e</sup> | no | yes, second degree<sup>g</sup> |  |
| yes | sMel | *D. melanogaster* | yes<sup>c</sup> (MSRO) | yes: wasp | yes<sup>f</sup> | yes, several sub-strains | not examined (strain discovered in 2005) | particles isolation, short-read sequencing |
| yes | sHy1 = sHyd1 | *D. hydei* | no | yes: wasp | no | yes, several sub-strains | yes, first degree<sup>g</sup> |  |
| yes | sNeo | *D. neotestacea* | no | yes: wasp and nematode | no | yes | not examined |  |
| unknown<sup>a</sup> | sEqu | *D. equinoxialis* | yes (ESRO) | unknown | no | no | yes, second degree<sup>g</sup> |  |
| unknown<sup>a</sup> | sPau | *D. paulistorum* | yes (PSRO) | unknown | no | no | yes, second degree<sup>g</sup> |  |

<sup>a</sup> these strains have not been studied since [56], and thus, their phylogenetic position has not been determined.
<sup>b</sup> the current common practice for *Spiroplasma* (and *Wolbachia*) is to use "s" for *Spiroplasma* (and "w" for *Wolbachia*) followed by the first three letters of the host species name.
<sup>c</sup> non-male-killing strains also occur, several have evolved within the lab, and have been used to elucidate the male-killing mechanism [e.g. 29, 120].
<sup>d</sup> the common practice was to use the first letter of the host species name followed by "SRO" or "SR", for Sex Ratio Organism
<sup>e</sup> First successful cell-free culture effort [121]; the second successful cell-free culture effort [122] was used as the basis for the description of *Spiroplasma poulsonii*, but to our knowledge was never replicated, and not found in any lab at present.
<sup>f</sup> this is the third successful cell-free liquid culture [66]; it has been replicated several times by the original authors. Our attempts to date have been unsuccessful (unpublished).
<sup>g</sup> see text for explanation; [24; and references therein, 43]
TEM = Transmission Electron Microscopy

*Spiroplasma* phage research was particularly active in the 1970’s [reviewed in 19, 41, 42-44], when electron micrographs of *Spiroplasma* cells (or their surrounding environment) that had been treated (e.g. exposed to broth culture or to particle extracts from a different *Spiroplasma* strain) or untreated, revealed the presence of viral-like particles of several distinct shapes. A subset of these particles, found in *Spiroplasma* cultivable strains (not associated with *Drosophila*), were propagated and further characterized [e.g. 45]. Filamentous or rod-shaped particles associated with non-lytic infections, were found protruding from the cell surface of numerous strains, including the *poulsonii* clade, as well as its closely related *citri* clade [42]. These particles are classified as plectroviruses, and contain circular ssDNA (7.5–8.5 kbp) in the virion, and dsDNA in the replicative form [46]; several (from the *citri* clade) have been sequenced (NCBI Acc. Nos. NC_003793, NC_001365, NC_009987, AJ969242, NC_001270). Sequences of plectroviral origin occur at many positions in the genomes of both *citri* and *poulsonii* clades [27, 47, 48, 49 and references therein]; a reason why complete chromosome assembly has been challenging. Two less common viral particles types (untailed isometric and long-tailed icosahedral) containing ssDNA are found exclusively in the *citri* clade [50, 51]. The fourth type of particle is a short-tailed icosahedron reported in the *citri* clade and in all the *Drosophila*-associated *Spiroplasma* strains known at the time [24, 42, 43, 45, 52-55]; all of which are known or suspected members of the *poulsonii* clade (four male-killers: NSRO, WSRO, ESRO, PSRO, and one non-male-killer: *s*Hyd1; see Table 1). These *poulsonii-*clade phages were characterized on the basis of particle morphology, nucleic acid content type, size, and/or phenotypic effects (i.e., evidence of *Spiroplasma* lysis and *Drosophila* sex ratio outcomes) of extract transfers among *Spiroplasma*-infected flies [24 and references therein, 56]. The first *Spiroplasma* strain from which a phage particle was isolated is NSRO (then thought to be a spirochaete); isolation was achieved without the need for “induction” by an outside source [“manifest” condition sensu 53]—hereafter termed “first-degree” extraction (Table 1). The only other *Spiroplasma* strain from which particles were isolated via the first-degree extraction is *s*Hyd1 [52]. Particles were isolated from the remaining *Drosophila*-associated strains following “induction”—i.e., exposure to a first-degree extraction [“latent” condition sensu 53]—hereafter termed “second-degree” extraction. A series of studies revealed that: (a) the particles of each strain infected, multiplied, and caused the lysis of at least one other strain; (b) the particles contained linear dsDNA genomes of 17, 21.8, or > 30 kbp, (circularly permuted in NSRO and *s*Hyd1; the only strains that produced sufficiently abundant particles to assess); and (c) NSRO harbored two different 21.8 kbp genomes [56]. To our knowledge, genome sequences have not been reported for any *Drosophila*-associated *Spiroplasma* short-tailed icosahedral particles containing dsDNA, or for the similarly shaped/sized dsDNA phages from the *citri* clade (genome size range: 16–21kbp), which include propagated lytic [45] and lysogenic [57] forms.

One caveat of the previous studies on *Drosophila*-associated *Spiroplasma* phage is that at that time, the existence of *Wolbachia* as a common endosymbiont of *Drosophila* (and many arthropods) was not known and therefore not tested. Of the native and non-native *Spiroplasma* hosts used in prior phage studies (see Table 1), at least *D. melanogaster, D. willistoni* and *D. paulistorum* but not *D. hydei*, are known to harbor *Wolbachia* [58, 59]. Therefore, it is possible that one or more of the phage attributed to *Spiroplasma* on the basis of morphology and genomic features by previous studies could instead represent *Wolbachia* phage WO [reported to have isometric heads ca. 40nm in diameter containing linear dsDNA ca. 20kbp; 60, 61].

Following a ∼4-decade hiatus, herein we re-launch research into phage particles of *Drosophila*-associated *Spiroplasma*. We isolated, visualized, extracted, and sequenced DNA (long and short reads) from phage-like particles. Our primary target was a representative of *S. poulsonii* strain NSRO, because NSRO is one of two strains from which past studies readily isolated phage particles via a “first-degree” extraction [43, 54]. The genome sequence of NSRO has not been published, so we were not able to compare the NSRO particle-derived contigs directly to the host genome. However, we used as a proxy for the host genome, the more recently discovered MSRO, native to *D. melanogaster*, which is the closest known relative of NSRO [62-64]. MSRO has been sequenced [several sub-strains; 27, 29, 65], and successfully cultured by one research group in cell-free liquid media [66] (see Table 1). To determine whether particles could be isolated from MSRO and assess the reproducibility of our isolation method, we performed isolation and sequencing (short reads only, due to much lower yield; see Results) of putative phage-like particles from one MSRO strain.

## Material and Methods

### Source of Biological Material

A *Drosophila melanogaster* Oregon R fly lab strain harboring *Spiroplasma poulsonii* NSRO [originally from D. nebulosa; 67] or MSRO-Br [originally from fly strain Red-42; 62] was maintained on Banana-Opuntia food at 25ºC (12:12 light:dark cycle). These two Oregon R lines infected with *Spiroplasma* were generated many generations ago by hemolymph transfer [e.g. as described in 67]. Both infected fly strains were maintained by mating to *Spiroplasma*-free Oregon R males. This *Spiroplasma*-free Oregon R fly strain served as a negative control in the phage isolation procedures. Adult flies were aged 1–2 weeks before phage extraction. Absence of *Wolbachia* infection was confirmed by PCR with *Wolbachia*-specific *wsp* primers [68, 69].

### Isolation of Phage-Like Particles

To enable the use of a large number of *Drosophila* individuals (i.e., > 3 grams), the PEG-based precipitation approach of van der Wilk et al. [70] was adapted as follows. Whole flies were homogenized with a mortar and pestle in 50ml of SM buffer (50 mM Tris–HCl, pH 7.5, 0.1 M NaCl, 10 mM MgSO_4_-7H_2_O), 1% (w/v) gelatin (VWR Cat. No. 97062-618) and 1μg/ml RNase A (Thermofisher Cat. No. EN0531; enzymatic activity: 5000 U/mg protein =100 Kunitz units/mg protein]. The homogenate was incubated at room temperature for 30 min and then centrifuged at 2000 x g to remove large debris. The supernatant was passed through a 70 μm filter (Millipore Cat. No. 28143-312) and incubated at 4ºC overnight. Our preliminary experiments indicated that thicker centrifugation bands were obtained if this incubation step was performed. Solid NaCl was added to final concentration of 1.0 M and incubated on ice for 1h. Solid PEG-6000 (ThermoFisher Cat. No. A17541.30) was then added to the solution to a final concentration of 10% (w/v) and gently mixed to dissolve. The mixture was incubated on ice for 1 h and then centrifuged at 11,000 x g for 10 min at 4ºC. The supernatant was decanted and discarded, and the pellet was resuspended in 8 ml of SM buffer by mixing for 10–20 min. An equal amount of chloroform was added to remove residual PEG. After gently mixing, the sample was centrifuged at 3,000 x g for 15 min at 4 ºC. The aqueous layer was collected and passed through a 0.45 μm filter (Acrodisk Cat. No. 28143-312). The sample was stored at 4ºC until ultra-centrifugation (i.e., for 24 to 48 h). The sample was subjected to an Optiprep (60% iodixanol) step gradient as previously described [71]. Each sample was layered over a prepared step iodixanol gradient of 10%, 30%, 35%, 40% and centrifuged at 209,500 x g at 4°C in a Beckman Coulter SW41 rotor for 2 h (Fig. 1a). Approximately 0.5ml were extracted from each interphase with a needle and stored at 4ºC until further use [i.e., Transmission Electron Microscopy (TEM) and/or DNA isolation]. These procedures were performed separately on NSRO-infected, MSRO-Br-infected, or uninfected flies (to serve as a *Spiroplasma*-free control). Additionally, to replicate the original findings in Oishi and Poulson [54], we isolated particles in a Cesium Chloride (CsCl) step gradient at a density of 1.48 g/cm^3^ for NSRO-infected and control fly homogenates only; these particles were only subjected to TEM.

**Fig 1.**
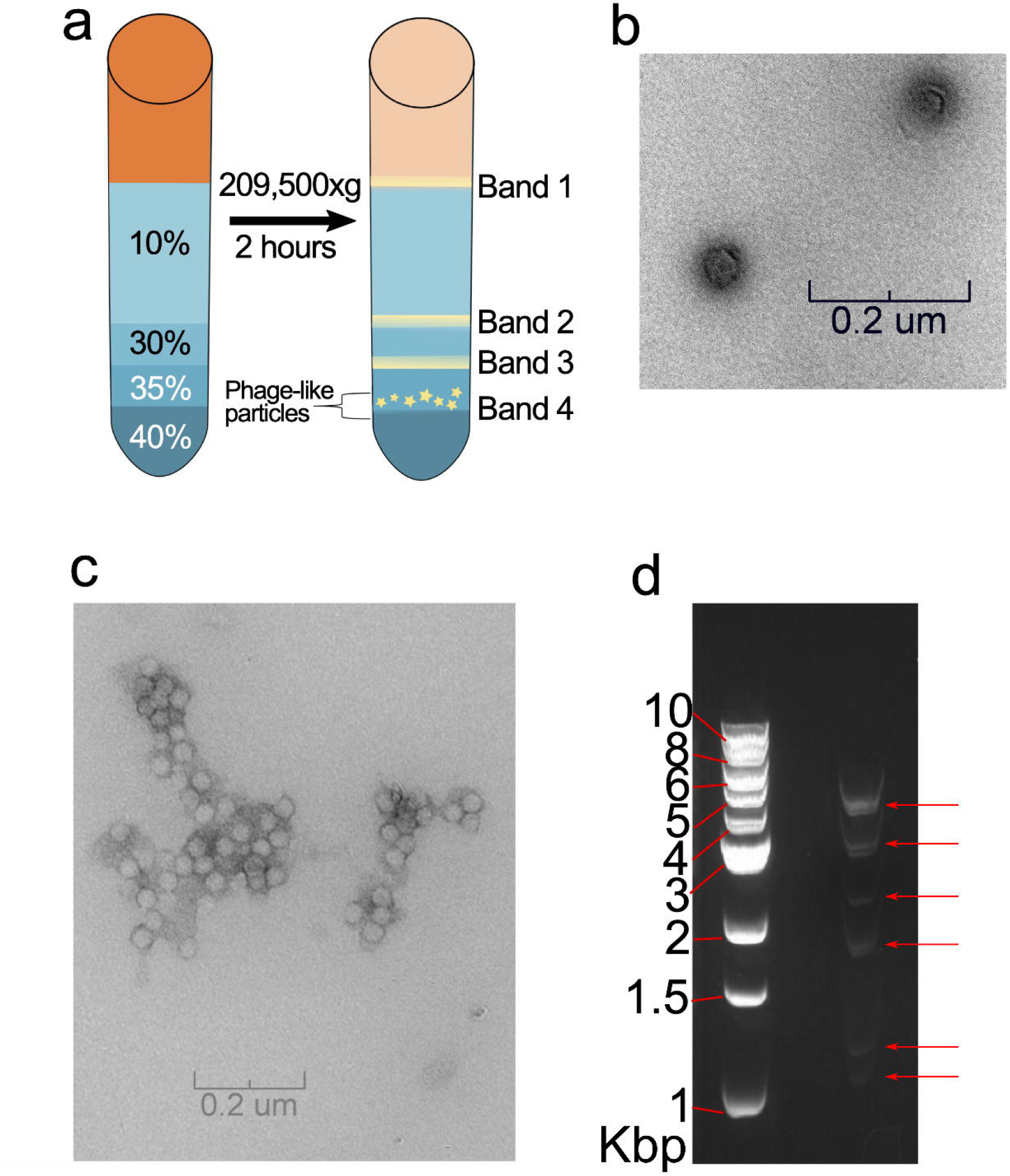
Isolation of Spiroplasma NSRO phage-like particles and their DNA. (a) Cartoon of the banding patterns obtained after Optiprep gradient centrifugation of the *Spiroplasma* NSRO extract aimed at isolating phage particles. Gradient layers (10, 30, 35, and 40% Iodixanol) are indicated on the left. Post-centrifugation (right), four bands (1–4) were visible. Extractions from the four bands were processed and viewed under Transmission Electron Microscopy (TEM). Only Band 4 (in the 35–40% interphase; indicated by stars) contained particles consistent with viral particles (shown in panel b). (b) TEM of negatively stained particles from band 4 in panel a. (c) TEM of negatively stained particles isolated from *Spiroplasma* NSRO with Cesium Chloride-based gradient at a density of at 1.48g/ml. (d) Gel electrophoresis of DNA extracted from the NSRO phage-like particles (right lane); visible bands are indicated with red arrow; an aliquot of this DNA isolation was subjected to Nanopore sequencing. Left Lane = 1 Kb DNA ladder (New England Biolabs Cat. No. N3232; band sizes indicated in Kbp).

### Transmission Electron Microscopy (TEM)

To search for phage-like particles, the interphase extracts of the NSRO strain were subjected to negative-stain TEM, following preparation by the Valentine method [72]. Interphase extracts were adsorbed to a carbon film, stained with 2% (w/v) uranyl acetate for 10–15 seconds, and placed on a copper grid (GT200T/Electron Microscopy Sciences). The grids were imaged at a magnification of 25,000x in a JEOL 1200 EX electron microscope (100 keV accelerating voltage) at the Texas A&M Microscopy and Imaging Center. The MSRO-Br extract was not subjected to TEM due to lack of sufficient sample.

### DNA Isolation from Particle Extracts

Only one interphase extract had a visible band and revealed phage-like particles under TEM (see Results). DNA was extracted from this interphase (both NSRO and MSRO-Br strains) with a Phage DNA isolation kit (Norgen Biotek Cat. No. 46800) according to the manufacturer’s protocol (elution volume = 50μl). DNA was quantified with a PicoGreen assay on a plate reader with a concentration of 100ng/μl. DNA was then cleaned with 0.9x AMPureXP beads (Beckman Coulter™ Agencourt Cat. No. A63880) to remove any residual EDTA, and resuspended in 24μl EB buffer (Tris HCl pH=8.5; Qiagen Cat. No. 19086).

### Library Preparation and Sequencing

Because the DNA amount was insufficient for the library preparation protocols for MinIon Nanopore (NSRO only) and Illumina MiSeq sequencing (NSRO and MSRO-Br), each DNA sample was separately combined with (“spiked into”) a larger DNA sample from an unrelated source. For the NSRO Nanopore library, we used DNA from the tephritid fruit fly *Anastrepha obliqua* to obtain ca. 10 μg of gDNA. Absence of *Spiroplasma* in the sample from *A. obliqua* was verified by PCR with *Spiroplasma*-specific primers targeting the 16S rRNA [16STF1 and 16STR1; 21]. The Nanopore SQK-LSK109 sequencing kit was used following the manufacturer’s recommendations (selected options: no DNA fragmentation; and use of HSB buffer to enrich for reads > 3 kbp). The Nanopore sequencing run consisted of a Spot-on flow cell Mk 1 R9 (FLO-MIN106), MinION Mk1B sequencer, and utilized the Nanopore sequencing software Minknow v2.1 (Oxford Nanopore). Basecalling was performed with Albacore v2.3.1 (Oxford Nanopore). Resulting reads were processed with PoreChop (https://github.com/rrwick/Porechop) to remove adapter sequences.

Two separate Illumina libraries were prepared: one for the NSRO sample and one for the MSRO-Br sample. Because the DNA amount from both samples was insufficient, prior to library preparation each of these samples was separately combined with (“spiked into”) much larger DNA amounts obtained from known and distinct phages harvested from bacterial cultures (i.e., *Vibrio, Staphylococcus*, and *Klebsiella*; see Results). The two Illumina libraries were prepared with TruSeq Nano low-throughput (LT) kit (Illumina), generating inserts with an average size of 580 bp. The Illumina libraries were sequenced with the MiSeq Platform (paired-end 150 bp). Hereafter, the total reads from the MSRO-Br Illumina sequencing will be referred to as the “MSRO-Br read set”, and likewise, the NSRO particle-derived reads will be referred to as “Illumina NSRO read set” and “Nanopore NSRO read set”.

### Initial assembly of Nanopore reads to retrieve putative *Spiroplasma*-derived contigs

The Nanopore NSRO read set was used alone for a preliminary assembly aimed at extracting the reads derived from the NSRO putative phage-like particles (as opposed to the DNA derived from the *Anastrepha obliqua* sample). The adapter-free fastq Nanopore reads were assembled with Canu v.1.7 [73] pipeline (genomeSize=2.0m, correctedErrorRate=0.105, corMinCoverage=0, corMaxEvidenceErate=0.15, corOutCoverage=1000; to maximize the number of reads utilized), then searched using blastn-discontiguous parameters to the NCBI nt database (January 2018). Three assembled contigs were identified as putative *Spiroplasma*-derived contigs based on an e-value of zero.

### Assessment of evidence of circular permutation in NSRO particle-derived contigs

One of the originally assembled putative *Spiroplasma* contigs was ∼ 31kbp, and contained flanking repeat regions (see Supporting Fig.S1); a pattern that is typically found in circularly permuted linear genomes [74]. To simplify downstream analyses, we removed one of the repeated regions to generate a “single copy” contig. This single-copy contig was used as a reference to re-map the original reads and generate a consensus. In addition, to determine if the original template molecules used to generate a contig were circularly permuted, we examined the pattern of read coverage in an artificial concatemer of the “single copy” contig (i.e., the “single copy” contig was tandemly repeated to generate a three-copy concatemer). The raw reads were then mapped to the concatemer to visualize mapping patterns. A gap in coverage would imply that all the template molecules share the same end(s), and thus, do not represent circularly permuted elements. The same process was repeated with a second contig that was originally 21kbp (see Supporting Figure S2). The third contig was removed from the assembly due to lack of coverage.

### Use of Illumina-derived reads to correct Nanopore-only contigs and to generate a hybrid assembly

To correct commonly encountered errors of Nanopore-only assemblies (i.e., insertions and deletions), the Illumina NSRO read set was aligned (10 iterations using the “map to reference” tool of Geneious v.11.1.2; Biomatters, Ltd.) to the remaining two single-copy-Nanopore contigs, and consensus contigs were generated (hereafter “Illumina-corrected-Canu” assembly).

We also implemented a hybrid de novo assembly using the Nanopore and Illumina reads. To simplify this assembly process, we first identified and discarded the Nanopore reads that did not map to the “Illumina-corrected-Canu” assembly, which were deemed of an origin other than *Spiroplasma*, such as the tephritid fly used to prepare this library. The mapping was achieved with Minimap2 (-ax map-ont --secondary=no; [75]). The retained “*Spiroplasma* only” Nanopore (long) reads and the quality-filtered “Illumina NSRO read set” were then subjected to a SPAdes v.3.12 hybrid assembly (33,55,77,107 kmer and default settings). The assembled contigs were then subjected to a Blast search [76, 77] using blastn-discontiguous parameters to the NCBI nt database (January 2018) to identify and retain putative *Spiroplasma* contigs with a maximum e-value of 1e-50 (hereafter, the “Hybrid Spades” assembly).

### Final NSRO contig assembly: reconciliation of “Illumina-corrected-Canu” and “Hybrid Spades” assemblies

To obtain the most accurate assembly possible, the “Illumina-corrected-Canu” and the “Hybrid Spades” assemblies were combined. This was done by aligning the “Hybrid Spades” assembly (which contained smaller contigs) to the “Illumina-corrected-Canu” assembly. “Hybrid Spades” contigs that aligned to the Canu reference with high identity (>99%), were considered to be part of, and merged with, the reference (“Illumina-corrected-Canu”) contig. The remaining “Hybrid Spades” contigs plus the all of “Illumina-corrected-Canu” contigs were considered as the final assembly. For the two largest contigs, which exhibited evidence of potential circular permutation, the origin was set so that they aligned to highly similar respective regions in the published *Spiroplasma* MSRO genomes (see Results).

Coverage and read lengths supporting the final NSRO particle-derived contigs were evaluated by mapping the “Nanopore NSRO read set” (with MiniMap2) and “Illumina NSRO read set” (with Geneious’ “map to reference” tool, with parameters: medium sensitivity; no iterations).

### Assembly of MSRO-Br particle-derived contigs (short reads only)

The MSRO-Br read set was assembled with SPAdes v.3.12 assembly (33,55,77,107 kmer and default settings). Akin to the procedures with the NSRO assembly, to identify the potential source of the assembled contigs, a blastn-discontiguous search was performed using the NCBI nt database (January 2018). Contigs with hits to *Spiroplasma* strains having an e-value of 1e-50 or less were regarded as derived from the MSRO-Br putative particles. Coverage of the final MSRO-Br particle-derived contigs was evaluated by mapping the “MSRO-Br read set” (with Geneious’ “map to reference” tool, with parameters: medium sensitivity, no iterations).

### ORF Annotation

Open reading frames (ORFs) from the NSRO and MSRO-Br assemblies were initially predicted and annotated using Prodigal [78], within the Prokka v.1.13 [79] pipeline with genetic code 4 (i.e., for *Mycoplasma*/*Spiroplasma*), which utilized a database containing all available NCBI *Spiroplasma* genomes (June 2018). Predicted ORFs at least 100bp long were retained, allowing for overlapping ORFs [80]. Non-coding RNA regions were annotated with Aragorn v.1.2.38 [81]. Assemblies were queried for potential prophage sequences using Phaster [82]. Additionally, translated ORFs were annotated with RASTtk v.2.0 [83; accessed June 2019] and searched manually by HHPred [84] and CD-Search [85] to assign functional annotations. Repeat regions were identified with the repeat finder tool of Geneious (parameters: at least 50bp long and 0% mismatches allowed). Signal peptide and transmembrane domains were assigned based on positive hits to the SignalP [86] and Phobius [87] databases. Similarly, putative transmembrane regions were annotated if positive hits were observed from both the TMHMM [88] and PHOBIUS [89] databases. IsFinder [90; accessed 27 June 2019] was used to identify possible transposable-element-associated proteins in the translated sequences with the IsFinder blastp database; a threshold e-value of 1E–40 was used to identify transposase families. Additionally, annotated genes, protein product names, and Interpro motifs names were visually inspected for terms associated with common toxins or virulence factors. The Geneious “Repeat finder” tool (parameters: at least 10bp long and 0% mismatches allowed) was used to manually identify inverted repeats flanking ORFs in assembled contigs (see Results).

### Comparison of particle-derived contigs to sequenced *Spiroplasma* genomes and phage

Ten genomes from *Spiroplasma poulsonii* and from the *citri* clade were analyzed for endogenous presence of our particle-derived contigs or parts of them, using a homology-based search with RepeatMasker v.4.0.9 [91]. Detected regions were filtered based on a minimum query coverage of 50%, and a Smith-Waterman (SW) score of less than 4.0%. Individual ORFs of our phage-like contigs were also searched with RepeatMasker in MSRO-Ug and MSRO-Br genomic assemblies to identify potential multicopy ORFs in the respective genomes (same minimum scores were used as for contig search).

To identify potential homology with previously reported sequences assigned to genomes of *Spiroplasma* viruses, we searched the NCBI nucleotide database (Date: 26 August 2022) for the terms “virus”, “phage”, “plectroviridae”, and “microviridae” in combination with the term “*Spiroplasma*”. The results were further filtered to retain only those in RefSeq and/or Species = “Viruses”, and to exclude those from metagenomic studies. We retained the following records: Plectroviridae (NCBI Acc. Nos. NC_009987, NC_003793,NC_001365, NC_001270, AJ969242) and Microviridae (NC_003438). In addition, a literature search revealed the existence of two *Spiroplasma citri* derived sequence fragments (U44405.1 and U28972.1) with “*Spiroplasma* phage SPV-1-like” features [92, 93]. We used Muscle v. 3.8.425 [94, 95] to attempt alignment of our particle-derived contigs with the above sequences.

### Assessment of expression of particle-derived contigs with available RNA sequencing datasets

To determine whether regions assembled in the particle-derived contigs show evidence of expression, we used publicly available RNA-seq data. An RNA sequencing dataset corresponding to *Spiroplasma* strain MSRO-Ug cultured in vitro and extracted from fly hemolymph was downloaded for NCBI GEO (GSE112290). Reads were trimmed and filtered with FASTP [default settings; 96]. Processed reads were then aligned with the Geneious RNA aligner (“medium sensitivity and span RNA introns” settings) to either the NSRO particle-or MSRO-Br particle-derived contigs. Alignments were then visually inspected with Geneious to qualitatively determine which of the ORFs in the particle-derived contigs were being expressed. Expression levels of entire contigs were computed using Reads per Kilobase Million (RPKM) method, except that gene length was replaced with the corresponding contig length. Expression was estimated by taking the number of reads aligned to each contig and dividing by the total number of sequenced reads. Then the resulting number was multiplied by a million to obtain a scaling factor to normalize for sequencing depth. The total number of reads aligned to a contig was then divided by both scaling factor and contig length in kilobases to obtain an RPKM-like value. Geneious was used to calculate raw reads and RPKMs of ORFs of NSRO or MSRO-Br particle-derived contigs.

## Results

### Isolation of phage-like particles

The iodixanol (Optiprep) step gradient of the NSRO sample resulted in a visible band at the interface of the 35–40% layers (Fig. 1a), consistent with the migration position for *Caudovirales* phages, which is similar in structure to previously described podoviridae-like *Spiroplasma* phage [71]. Such a band was barely visible in the MSRO-Br sample, and it was not visible in the control samples of *Spiroplasma*-free fly homogenates. Extractions generally yielded a more visible band in the Optiprep gradient when flies were aged for at least two weeks, and the filtered (0.45 μm) homogenate was allowed to incubate for at least 24 h (results not shown). The Cesium Chloride (CsCl) gradient yielded a band at density of 1.48 g/cm^3^ (consistent with [54]). TEM of the Optiprep 35–40% interphase and the CsCl band, revealed tail-less, icosahedral particles measuring ca. 40 nm across (Fig. 1b,c; respectively). DNA extracted from Optiprep 35–40% interface from the NSRO sample produced multiple bands when run on an agarose gel, visible at approximately 1–1.5, 2, 2–3, 3–4, and 5 kbp (indicated with arrows in Fig. 1d). TEM and gel electrophoresis were not performed on the MSRO-Br sample due to its low yield, which was subjected to Illumina DNA sequencing only.

### Assembly of NSRO and MSRO-Br particle-derived contigs

From the NSRO sample, three contigs with blast hits to *Spiroplasma* were assembled with Canu, but one was discarded due to low read coverage (7X coverage; only one read > 15kbp) and close sequence similarity to a higher coverage contig, resulting in two remaining contigs of size 31 kbp and 21 kbp. However, the presence of 12 kbp- and 1.3 kbp-long terminal repeats (oriented in the same direction) at their ends, respectively, and the raw read mapping pattern observed (Supporting Figs. S1 and S2), suggested that these original contigs contained assembly artifacts. These contigs were therefore adjusted to where a single repeat region was removed from each, leaving “single copy” contigs, resulting in two contigs ∼19 kbp-long each. These contigs were then corrected with Illumina reads. The resulting “Illumina-corrected-Canu” contigs were then reconciled with the SPAdes Hybrid assembly.

Four contigs from the reconciliation “Final” assembly were recovered. The two aforementioned ∼19 kbp contigs (hereafter referred to as “NSRO-P1” and “NSRO-P2”) were not evident in the gel of DNA isolated from the particle extract (Fig. 1d). The sizes of the remaining two contigs (Table 2) were consistent with the DNA bands observed in the gel. The two other contigs, hereafter Contig 3 and “NSRO-IS5”, were 3.8 and 1.2 kbp, respectively.

**Table 2.** Characteristics of the contigs derived from the putative phage particles with sufficient read coverage.

| Contig Name | Length<br>bp | GC<br>content<br>% | Number<br>of ORFs | Illumina Miseq<br>reads mapped<br>(coverage) | Nanopore Minion reads<br>mapped (coverage) |
| --- | --- | --- | --- | --- | --- |
| **NSRO** |  |  |  |  |  |
| NSRO-P1 | 18,836 | 24.0 | 23 | 1,795 (13.7) | 400 (51.4) |
| NSRO-P2 | 19,169 | 24.2 | 24 | 1,163 (8.79) | 268 (28.2) |
| 3 | 3,805 | 24.8 | 3 | 121 (4.33) | 2 (0.55) |
| NSRO-IS5 | 1266 | 21.6 | 2 | 55 (6.060) | 0 (0) |
| **MSRO-BR** |  |  |  |  |  |
| MSRO-BR-P1 | 19,700 | 24.9 | 22 | 1,298 (9) | NA |
| MSRO-BR_IS5 | 1,106 | 20.7 | 2 | 106 (5) | NA |

From the MSRO-Br sample, for which only Illumina reads were generated, we obtained six contigs with high homology to *Spiroplasma*, based on blastn, of which only two had adequate coverage: a 19.7 kbp contig (hereafter referred to as “MSRO-Br-P1”) along with a 1.1 kbp contig (hereafter “MSRO-Br-IS5”; Table 2). The other four contigs were of lengths shorter than 500 bp and considerably lower coverage (See Supporting Dataset S1).

The GC content range of NSRO-P1, NSRO-P2, Contig 3, and MSRO-Br-P1 was 24.0–24.9% (Table 2), whereas the GC content of NSRO-IS5 and MSRO-Br-IS5 was lower (21.6 and 20.7%, respectively). These values are similar to the GC content of the Plectroviridae class of *Spiroplasma* bacteriophages (∼23%), and the complete MSRO-Ug genome of 26.5% [65], but substantially lower than the Microviridae class (32.1%).

### Coverage and size distribution of reads mapped to the particle-derived contigs

400 Nanopore reads mapped to NSRO-P1 (not shown), of which 15 were at least 16 kbp, and none were greater than 22 kbp (Fig. 2a), suggesting that if this represents a phage genome, its size is less than 22 kbp (reads longer than 22 kbp were obtained from this sequencing library, but could be attributed to *A. obliqua* or its associated microbes; not shown). The average, maximum and minimum coverage of the Nanopore reads was respectively 52, 72, and 32X. The manner in which the longest Nanopore raw reads mapped to the artificial three-copy concatemer version of NSRO-P1, with no gaps in coverage for the middle copy, or in its 5’ and 3’ junctions with the adjacent copies (Fig. 3a), is consistent with a circularly permuted genome and a headful packaging mechanism [97]. 1,795 Illumina reads aligned to the NSRO-P1 single-copy contig (13.7X coverage; Table 2), of which 18 pairs had the following pattern consistent with circular permutation: one member of the pair mapped to one end of the contig whereas the other mapped to the other end (not shown).

**Fig. 2.**
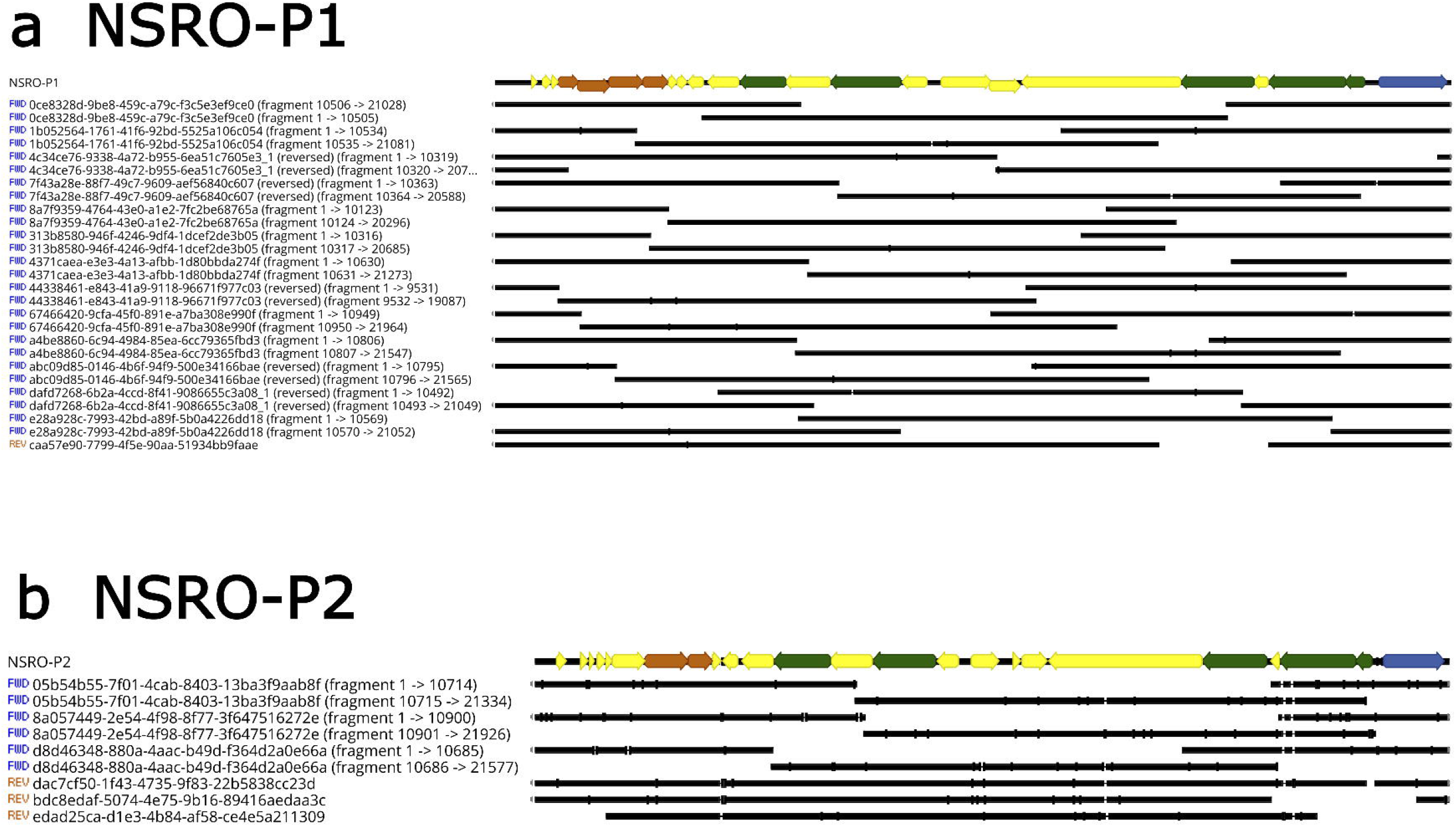
Mapping results of the trimmed long reads (black) to the NSRO-P1 and NSRO-P2 contigs. (a) The 14 longest reads (16 kbp to 21.9 kbp; avg 20 kbp) mapped to NSRO-P1 are shown. (b) The 7 longest reads mapped to NSRO-P2 (14.852 kbp to 21.9 kbp; avg 19 kbp) are shown. Light gray boxes in reads represent trimmed regions. Thin lines represent gaps in alignment of the reads to the reference. Annotated ORFs (colored arrows) are shown for reference and described in detail in Figure 6.

**Fig. 3.**
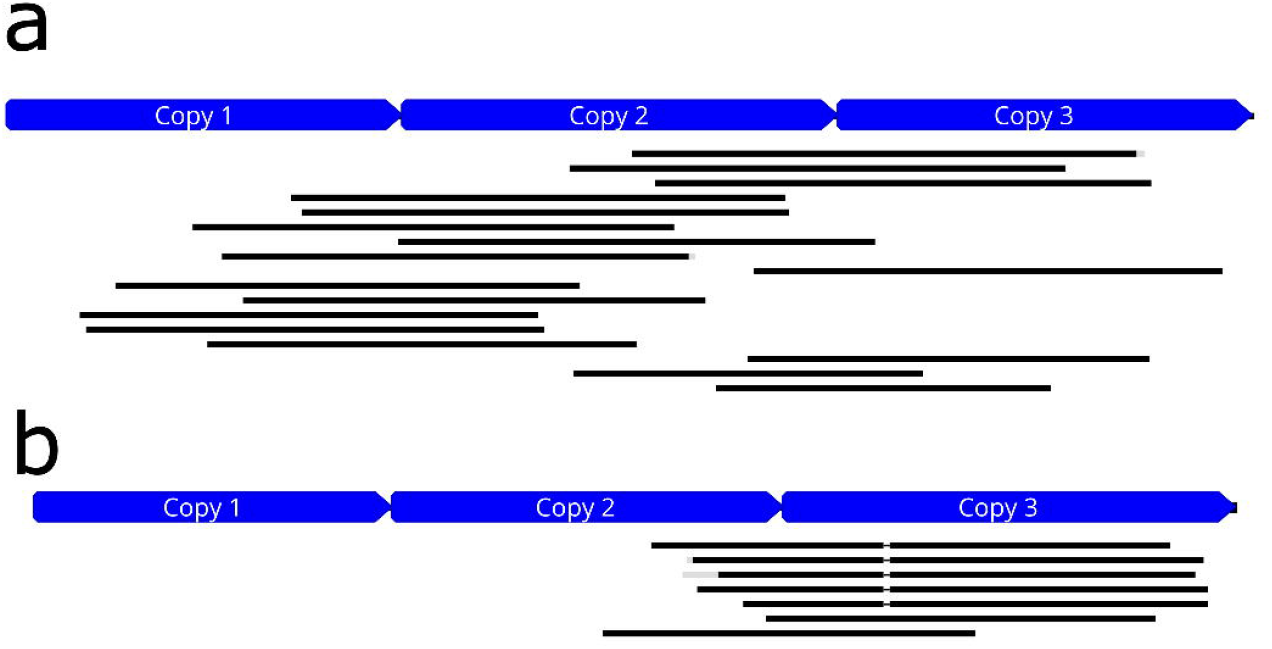
Mapping patterns of the longest Nanopore reads to artificial three-copy concatemers of NSRO-P1 (a) and NSRO-P2 (b) are consistent with a circularly permuted genome and a headful packaging mechanism. The raw long reads are shown in black. Gray outline boxes in reads represent trimmed regions (i.e., different from reference).

268 Nanopore reads mapped to NSRO-P2 (not shown) with an average, maximum and minimum coverage of 28.2, 42, 16X respectively. Seven Nanopore reads were > 14 kbp long and their mapping to the single-copy contig is shown in Fig. 2b. Mapping of these seven Nanopore reads to the artificial three-copy concatemer (Fig. 3b), also suggests circular permutation and headful packaging. 1,163 Illumina reads aligned to the NSRO-P2 genome (8.679X coverage), of which 10 pairs had each member mapped to opposite ends of the contig (not shown), consistent with circular permutation.

Read coverage for the Contigs 3 and NSRO-IS5 was lower (Table 2). Two Nanopore and 121 Illumina reads mapped to Contig 3 (Illumina coverage = 4.33X). Zero Nanopore and 55 Illumina reads mapped to NSRO-IS5 (Illumina coverage = 6X). Low to no coverage by Nanopore reads was expected because the library preparation procedure was aimed at enrichment of > 3kb fragments).

Coverage of MSRO-Br-P1 ranged from 1–31 Illumina reads with an average of 9 reads for a total of 1,298 reads (Table 2). Coverage of MSRO-Br-IS5 ranged from 1-10 reads with an average of 5 reads for a total of 106 reads.

### Functional annotation of particle-derived contigs

The final 18.8 kbp contig of NSRO-P1 contained 23 predicted protein-coding genes ranging from 41 to 1067 aa (Supporting Dataset S2; Fig. 4a). The contig resembles a partial phage genome, with large and small terminase subunits, portal, head-tail connector and capsid proteins identifiable by conserved domain or HHPred searches. In addition to these structural proteins, genes encoding phage-like DNA replication functions were also identified, including a single-stranded DNA binding protein, a potential replicative helicase/nuclease, a RecT-like recombinase and a Holliday junction resolvase. Two ORFs, peg.18 and peg.21 coded for hypothetical proteins with a bacterial signal motif and a eukaryotic signal motif, respectively. Additionally, three ORFs were predicted to contain transmembrane domains, peg.16 with three transmembrane domains, and peg.15 and peg.22 each with one transmembrane domain. This contig does not appear to be a complete phage genome, however, as it is notably missing recognizable regulatory switches, tail structural proteins, lysis genes, and an integrase that would be responsible for insertion and excision of the element. NSRO-P2, 19.1 kbp in length, contained 24 ORFs (size range 38– 1073 aa; Fig. 4b), many of which are shared with NSRO-P1. However, NSRO-P2 uniquely encodes an ORF (peg.25) that was assigned a putative function of prophage-associated RecT ssDNA binding protein (peg.35; Uniprot ID: PF03837). Two ORFs in NSRO-P2 coded for transmembrane proteins motifs, peg.38 (three motifs) and peg.39 (one motif), which are identical to peg.15 and peg.16 in NSRO-P1, respectively.

**Fig. 4.**
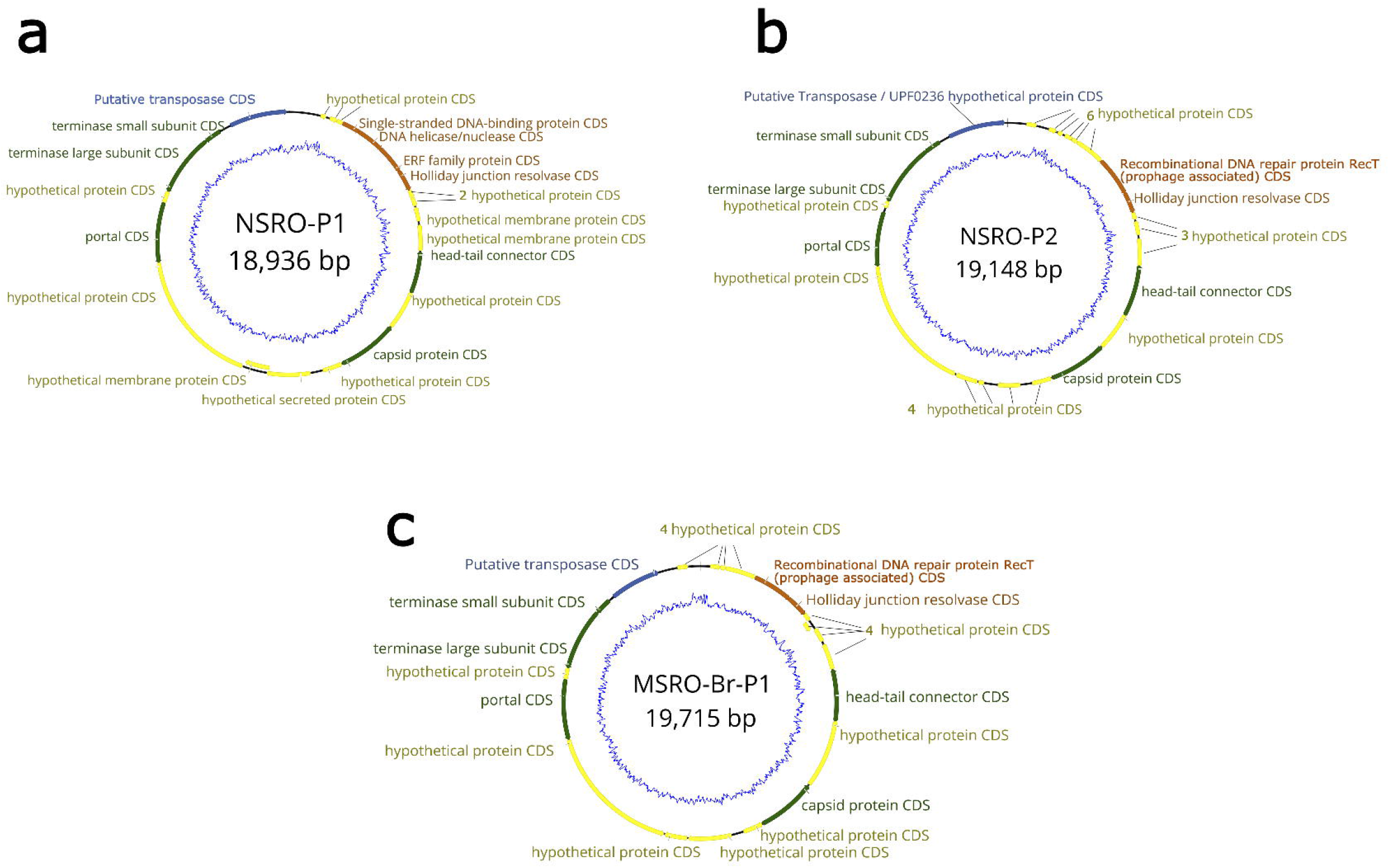
Annotation of phage-like particle-derived genomes. Open reading frames (ORFs) are indicated with colored block arrows. Yellow = hypothetical CDS with no homology to known proteins with known functions. Blue graphs depict the GC content at a 15 bp sliding window. (**a**) NSRO-P1, **(b**) NSRO-P2, (c) MSRO-Br-P1 contained annotated ORFs, Blue = transposase proteins. Green = capsid structure proteins. Orange = DNA binding/recombination proteins.

The entirety of Contig 3 (3.8 kbp) mapped with 98.7% similarity to a region (1,436,327–1,440,042) of the *Spiroplasma* MSRO-Ug assembly GCF_000820525.2_SMSRO_2016_genomic (as well as to closely related strains; see below). This region contains three ORFs, of which one (peg.3 in our contig) corresponds the MSRO-Ug’s Ribosome Inactivating Protein 2 (RIP2) gene containing a signal peptide, whose RIP-function has been confirmed experimentally [32]. Another ORF (peg.2 in our contig) also has a signal peptide motif and is found in NSRO-P1 (=peg.22), NSRO-P2 (=peg.46), MSRO-P1 (=peg.8; see below). NSRO-IS5 contained two ORFs (peg.4 and peg.5). The former has a DDE superfamily endonuclease, and the latter has a helix-turn-helix DNA-binding motif that is common among transposable elements (based on interpro: IPR027805). IsFinder indicated that both peg.4 and peg.5 had homology to transposable elements of the family IS5, group IS2. Additionally, using the Geneious repeat finder, we identified flanking inverted repeats (12bp long, 5’-AAAATATAATTT-’3) in NSRO-IS5. According to the NCBI Reference Sequence (WP_105628893.1), peg.4 and peg.5 undergo ribosomal slippage to produce a larger combined single protein composed of both a DDE superfamily endonuclease and a helix-turn-helix DNA-binding motif. A transmembrane motif also exists in the protein sequence.

MSRO-Br-P1 has 22 predicted ORFs (ranging from 45-1,071 aa) with 7 ORFs containing protein with known motifs according to InterPro, Phobius or SignalP (Fig. 4c). The contig contains genes with large and small terminase subunits, portal, head-tail connector and capsid proteins identifiable by conserved domain or HHPred searches. In addition, a RecT-like recombinase and a Holliday junction resolvase were identified (Supporting Dataset S2). MSRO-Br-IS5 was identical in annotation to NSRO-IS5 consisting of two ORFs and the observed inverted repeats (Supporting Dataset S2).

### Similarity among the particle-derived contigs from NSRO and MSRO-Br

MSRO-Br-P1 was aligned to both NSRO-P1 and NSRO-P2 with the Geneious Aligner (Global alignment with free end gaps, Cost Matrix: 65% similarity, Gap Open Penalty: 12, Gap Extension Penalty: 3). MSRO-Br-P1 shares 78% and 81% identity with NSRO-P1 and NSRO-P2, respectively (Supporting Fig. S3A). Interestingly, the recT-like recombinase of MSRO-Br-P1 is similar to the NSRO-P2 annotated recT gene (94.8%). Unlike NSRO-P1, a single-stranded DNA-binding protein or DNA helicase/nuclease protein was not annotated in MSRO-Br-P1. MSRO-Br-IS5 is identical to the NSRO-IS5 except on the terminal ends of the contigs (Supporting Fig. S3B).

### Comparison of NSRO and MSRO-Br particle-derived contigs to available sequenced genomes of *Spiroplasma* phage

Attempts to align the NSRO and MSRO-Br particle-derived contigs to previously described plectroviral phage and other phages from *Spiroplasma* revealed substantial homology to only the ∼9.6 kbp *S. citri* sequence U44405.1, whose pairwise identity to NSRO-P1, NSRO-P2, and MSRO-Br-P1 was 74.9, 74.1, and 76.5%, respectively (Fig. S4). U44405.1 is a fragment that was found to be deleted in a strain of *S. citri* that lost the ability to be transmitted from its insect to its plant host, and was hypothesized to contain a transposase and one or more adhesion proteins with the following molecular weights (P58, P12, P54, P123, P18; [92]). The ORFs annotated in U44405 share substantial synteny and predicted CDS boundaries, with those in our phage-like contigs. P58 is a membrane protein [98] belonging to a multigene family [99]. Whether any of the proteins encoded by U44405.1 are involved in insect transmissibility has not been further investigated [see 100]. In accordance with their homology to our phage-like contigs, HHPred annotated P58 as “large terminase subunit”, P54 as “portal”, and “transposase” as “transposase” (Fig. S4). HHPred did not reveal annotation terms for any other ORFs in U44405.1.

A genome assembly of *Spiroplasma poulsonii* strains NSRO is not currently available to search for evidence of the presence of our particle-derived contigs within the bacterial chromosome, and thus confirm them as originating from the *Spiroplasma* strain (e.g. if they are lysogenic). Nonetheless, we searched for regions homologous to the *Spiroplasma* particle-derived contigs in the main chromosomal contigs and associated plasmids of the currently available genomes of the members of the *poulsonii* and *citri* clades of *Spiroplasma* (Table 3). For the *poulsonii* clade, we included *S. poulsonii* Neo (associated with *D. neotestacea*), and four sub-strains of MSRO, three of which are derived from “MSRO-Ug”; a sub-strain of MSRO originally found in *D. melanogaster* from Uganda [101]. The fourth substrain is MSRO-Br, from which MSRO-P1 was extracted. Based on sequences from several genes, *S. poulsonii* MSRO and NSRO are each other’s closest relatives [62-64]. We found no regions of homology (criteria: minimum query coverage 50% and a Smith-Waterman score < 4) in the currently available *citri* clade genome assemblies; none of these *citri* clade strains are associated with *Drosophila*.

**Table 3.** Summary of the results from the search for regions with similarity to the NSRO and MSRO-Br particle-derived contigs in *Spiroplasma* genome assemblies from the *poulsonii* and *citri* clades. Only regions with high homology (Smith-Waterman <4.0 and > 50% length of original contig) were counted as complete regions or remnants of the NSRO putative phage contigs. Detailed output available in Supporting Dataset S6. All listed genomes are considered “complete” genomes except for Neo and MSRO-Br. MK = male killing. *S. poulsonii* belong to the *poulsonii* clade, whereas all other belong to the *citri* clade.

| GenBank assembly accession number | Phenotype | Species, strain, and chromosome, contig, or plasmid ID [size in kbp] | NSRO-P1 (18.8 kpb) | NSRO-P2 (19.2 kbp) | NSRO Contig 3 (3.8 kbp) | IS5 (1.26 kbp) | MSRO-Br-P1 (19.71 kbp) |
| --- | --- | --- | --- | --- | --- | --- | --- |
| GCA_000820525.2 | MK,Cultured | *S. poulsonii* MSRO-Masson (JTLV02000001) [1,800] | 3 | 5 | 1 | 236 | 5 |
|  |  | *S. poulsonii* MSRO-Masson Plasmid (JTLV02000002) [36.2] | 0 | 0 | 0 | 8 | 0 |
| GCA_003124105.1 | Non-MK | *S. poulsonii* MSRO-SE (NZ_PETF01000001.1) [1,800] | 3 | 6 | 1 | 237 | 7 |
|  |  | *S. poulsonii* MSRO-SE Plasmid (NZ_PETF01000002) [55] | 0 | 0 | 0 | 8 | 0 |

| GenBank assembly accession number | Phenotype | Species, strain, and chromosome, contig, or plasmid ID [size in kbp] | NSRO-P1 (18.8 kbp) | NSRO-P2 (19.2 kbp) | NSRO Contig 3 (3.8 kbp) | IS5 (1.26 kbp) | MSRO-Br-P1 (19.71 kbp) |
| --- | --- | --- | --- | --- | --- | --- | --- |
| GCA_003124115.1 | MK | *S. poulsonii* MSRO-H99 (NZ_PETG01000001.1) [1,400] | 2 | 4 | 0 | 174 | 7 |
|  |  | *S. poulsonii* MSRO-H99 Plasmid (NZ_PETG01000004) [87] | 0 | 0 | 0 | 19 | 0 |
|  |  | *S. poulsonii* MSRO-H99 Artifact (NZ_PETG01000003) [257] | 0 | 4 | 1 | 0 | 2 |
| GCA_019295385.1 | MK | *S. poulsonii* MSRO-Br (Red 42) | 0 | 0 | 1 | 3 | 1 |
| GCA_003989055.1 | Non-MK | *S. poulsonii* Neo | 0 | 0 | 0 | 265 | 0 |
| GCA_003339775.1 | Plant pathogen | *S. phoeniceum* strain:p40 | 0 | 0 | 0 | 0 | 0 |
| GCF_000236085.2 | Bee pathogen | *S. melliferum* | 0 | 0 | 0 | 0 | 0 |
| GCF_000328865.1 | Bee pathogen | *S. melliferum* | 0 | 0 | 0 | 0 | 0 |
| GCF_001274875.1 | Corn pathogen | *S. kunkelii* CR2-3x | 0 | 0 | 0 | 0 | 0 |
| GCF_001886855.1 | Plant pathogen | *S. citri* | 0 | 0 | 0 | 0 | 0 |

Contig NSRO-P1 mapped with high homology to 2–3 regions in the main chromosomal contigs of the three MSRO-Ug genomes (Table 3; Supporting Dataset S6). Similarly, NSRO-P2 mapped with high homology to 3–5 regions of the three MSRO-Ug genomes. The RIP2-containing Contig 3 mapped with high homology (i.e., 98.7% similarity) to a single location in the chromosome of two MSRO substrains (MSRO-Masson GCA_000820525.2 and MSRO-SE GCA_003124105.1), and at a single location of a contig deemed an extrachromosomal element (or assembly artifact) of MSRO-H99 (GCA_003124115.1). Contig 3 also mapped to a single contig of the MSRO-Br assembly (contig VIGU01000005) with 98.6% similarity. As mentioned above, the RIP2-encoding region of Contig 3 had 100% match to the corresponding regions in the four MSRO assemblies. NSRO-P1, NSRO-P2, and Contig 3 had no matches fulfilling our criteria in the other six *Spiroplasma* genomes examined.

NSRO-IS5 (and MSRO-Br-IS5; its identical counterpart in MSRO), or its remnants, were found in numerous locations of all four of the *poulsonii*-clade genomes examined (Table 3): found at 174–237 chromosomal and 8–19 plasmid locations of the three MSRO-Ug genomes; and at 265 locations in the genome assembly of *S. poulsonii* Neo. The IS5 contig was only found at 3 locations in the MSRO-Br draft genome, which is highly fragmented.

Incomplete parts of the MSRO-Br-P1 contig were found in the three MSRO assemblies (Table 3). The full MSRO-Br-P1 contig was found in the MSRO-Br whole genome assembly with 85% percent identity to the circular VIGU01000004 (18.9 kbp) contig. Additionally, MSRO-Br-P1 was found in 5–7 different regions of each MSRO-Ug assembly, with the largest regions being ∼12,000bp long. Interestingly, parts of the MSRO-Br-P1 contig were found in 13 regions belonging to putative assembly artifacts in the MSRO-JTV (GCA_000820525.2) assembly, and 6 regions in putative assembly artifacts in MSRO-H99 assembly (GCA_003124105.1).

### Assessment of contamination by DNA from sources other than *Spiroplasma*

Due to the low amount of DNA obtained from the extractions targeting phage-like particles, our DNA extracts were combined with (“spiked into”) a different sample that contained a much larger amount of DNA. In Supporting Dataset S3, we report the top blast hits of all contigs obtained as a result of our assemblies using all of the reads. For the Illumina MiSeq run (Index 4), which contained DNA from *Staphylococcus* and *Klebsiella* phages with our NSRO particle extraction, we obtained large contigs whose top blast hits were the known phage. Numerous contigs whose top blast hit was *Klebsiella* or other Proteobacteria were also recovered.

For the Nanopore run, the only contigs whose top blast hit was *Spiroplasma* were the ones we assigned to *Spiroplasma* NSRO particles. As expected from a library prep containing a predominance of DNA from *Wolbachia*-infected *A. obliqua*, numerous other contigs had blast hits to *Anastrepha, Wolbachia*, and Proteobacteria (not reported).

For the Illumina MiSeq run (Index 1), which contained DNA from *Vibrio, Staphylococcus* and *Klebsiella* phages, combined with our MSRO-Br particle extraction, we obtained large contigs whose top blast hits were these phage. In addition to the contigs that we assigned to *Spiroplasma* MSRO particles, we only detected four short contigs (259 to 468 bp) whose top blast hit was *Spiroplasma* (very low coverage). A few contigs whose top blast hit was Proteobacteria were also recovered, as well as a few short contigs (<400bp) whose top blast hit was *Homo sapiens*.

### Representation in RNA-seq datasets of the particle-derived contigs

To determine if there is evidence of expression (at the mRNA level) of the ORFs annotated in the particle-derived contigs, we used currently available RNA sequencing datasets from *Spiroplasma poulsonii* MSRO-Ug. These datasets were derived from fly hemolymph and from in vitro culture (NCBI PRJNA445479; GEO: GSE112290). We decided to exclude the RNAseq dataset of MSRO-Br [38], because it was based on whole insect extractions and the yield of *Spiroplasma* reads was relatively low. Based on Reads per Kilobase Million (RPKM), at the contig level, NSRO-P1 had the lowest expression level (average 16–17; Table 4; Supporting Dataset S4). At the ORF level, NSRO-P1 also had the least representation with only 18 out of the 23 ORFs having at least 5 reads per replicate (in culture; Table 4; Supporting Dataset S5). Higher RPKMs were observed at the contig level for NSRO-P2 (50 in culture; 31 in hemolymph) and MSRO-P1 (57 in culture; 42 in hemolymph). At the ORF level, at least 21 out of 24 ORFs of NSRO-P2 had at least 5 reads mapped per replicate, but small regions of ORFs lacked coverage. For MSRO-P1, at least 21 out of 22 ORFs had coverage of at least 5 reads. Regions lacking coverage by RNA-seq reads may reflect absence of template DNA (chromosomal or extrachromosomal) in MSRO-Ug, or lack of expression despite presence of template DNA (e.g. because it is not a functional gene or because expression of the functional gene does not occur under those experimental conditions). At the contig level, IS5 had the highest RPKM (294 in culture; 119 in hemolymph), and both ORFs were covered by at least 5 reads. Finally, NSRO Contig 3 (the one containing the RIP2 gene) had higher RPKM in hemolymph (43.5) than in culture (38.7) of MSRO-Ug, as reported by Masson et al. [66].

**Table 4.**
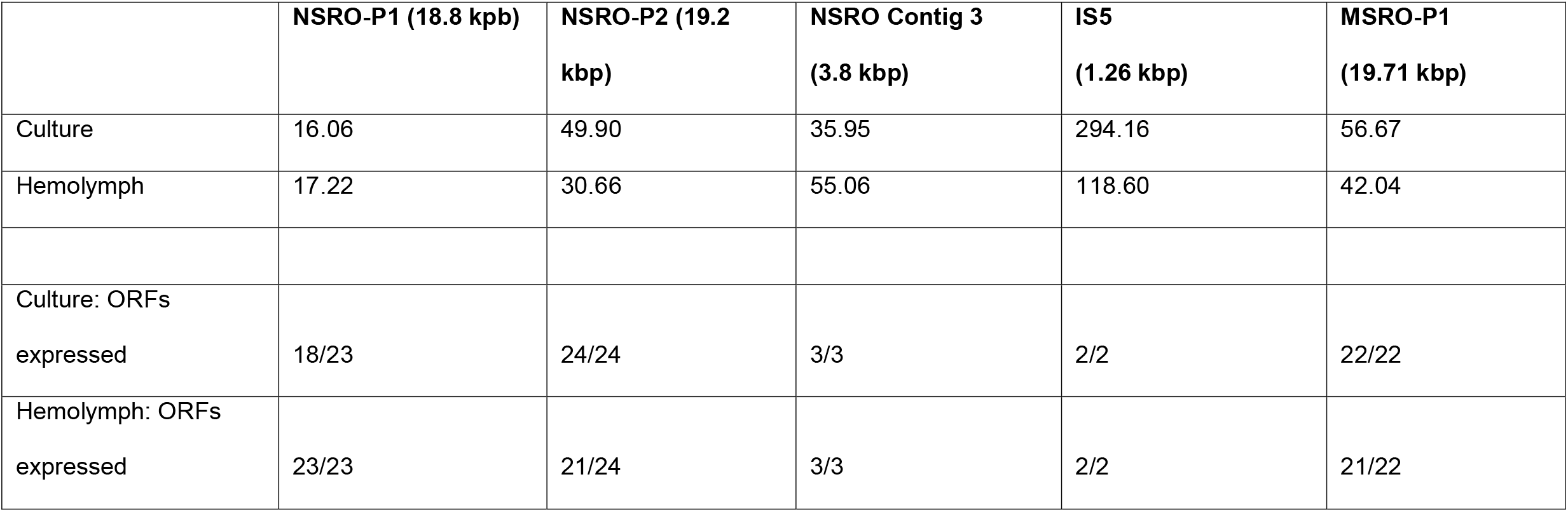
Analysis of mRNA expression levels of particle-derived contigs, by published *Spiroplasma poulsonii* MSRO-Ug RNA-seq datasets. RNA-seq reads were aligned to the five particle-derived contigs (from both NSRO and MSRO) to estimate expression at the contig level (Reads per Kilobase Million; RPKM). The numerator in “ORFs expressed” rows indicates ORFs that had at least five reads mapped per replicate. Detailed output by contig and ORF is provided in Supporting Dataset S4.

|  | **NSRO-P1 (18.8 kbp)** | **NSRO-P2 (19.2 kbp)** | **NSRO Contig 3 (3.8 kbp)** | **IS5 (1.26 kbp)** | **MSRO-P1 (19.71 kbp)** |
| --- | --- | --- | --- | --- | --- |
| Culture | 16.06 | 49.90 | 35.95 | 294.16 | 56.67 |
| Hemolymph | 17.22 | 30.66 | 55.06 | 118.60 | 42.04 |
| Culture: ORFs expressed | 18/23 | 24/24 | 3/3 | 2/2 | 22/22 |
| Hemolymph: ORFs expressed | 23/23 | 21/24 | 3/3 | 2/2 | 21/22 |

### Coverage of RIP2 contig in Illumina DNAseq datasets not aimed at particles

The recovery of RIP2 (from NSRO contig 3) in our particle-derived contigs is interesting because of its ribosome depurination activity, the potential role of RIPs in causing death of parasitic wasps, as well as of *Drosophila* embryos and old adults, RIP2 expression patterns (relatively high expression, and differential expression among conditions), and the fact that RIP2 is regarded as a single-copy gene in all (MSRO) assemblies to date [27, 32, 36, 38, 39, 66]. We determined whether its coverage by MSRO Illumina DNAseq datasets not aimed at particles (e.g. those used to generate genome assemblies) is consistent with a single-copy gene. We found that its coverage in MSRO-Ug (SAMN15339386) was consistent a single-copy gene, but its coverage in MSRO-Br (SAMN10488582) suggested ∼14x copies compared to other presumably single-copy genes (e.g. *dnaA*; Table 5).

**Table 5.**
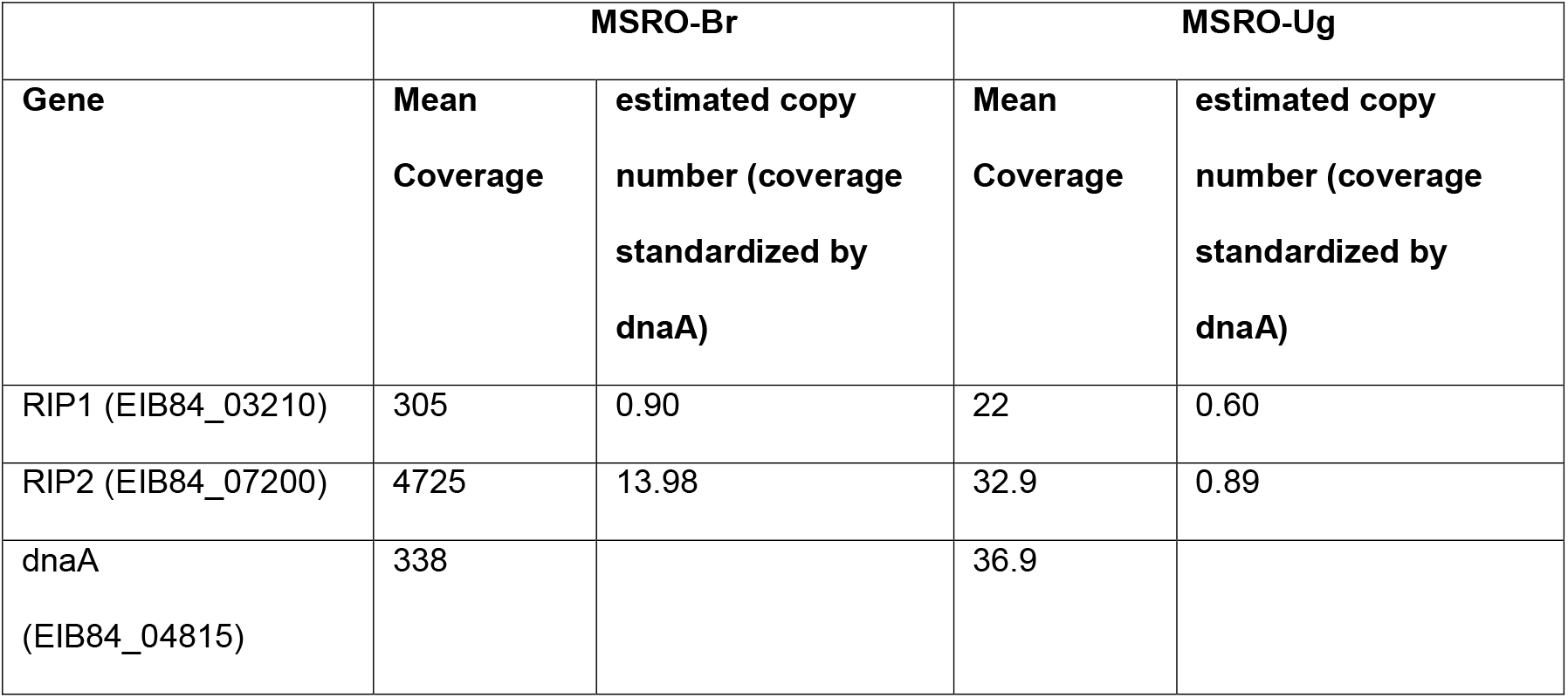
Comparison on inferred copy numbers of RIP1 and RIP2 genes in *Spiroplasma poulsonii* substrains MSRO-Br (NCBI Acc. Nos.: PRJNA507275; SAMN10488582) and MSRO-Ug (NCBI Acc. Nos.: PRJNA640980; SAMN15339386). Mean coverage of RIP1, RIP2, and dnaA (presumably a single-copy gene). Locus tags in MSRO-Br genome (VIGU00000000.1) are shown in parenthesis. Mean coverage was estimated in Geneious by examining the output of reads mapped to the three genes by Bowtie2 [123] with default parameters. Within each substrain, mean coverages of RIP1 and RIP2 were divided by mean coverage of dnaA to estimate their copy numbers.

|  | **MSRO-Br** |  | **MSRO-Ug** |  |
| --- | --- | --- | --- | --- |
| **Gene** | **Mean Coverage** | **estimated copy number (coverage standardized by dnaA)** | **Mean Coverage** | **estimated copy number (coverage standardized by dnaA)** |
| RIP1 (EIB84_03210) | 305 | 0.90 | 22 | 0.60 |
| RIP2 (EIB84_07200) | 4725 | 13.98 | 32.9 | 0.89 |
| dnaA<br>(EIB84_04815) | 338 |  | 36.9 |  |

## Discussion

The main goal of this study was to assemble the genome of phages described in several *Drosophila*-associated *Spiroplasma* strains before the advent of DNA sequencing. Several of our findings suggest that our density step gradient isolations recovered particles that may correspond to the previously described short-tailed dsDNA phages from *Drosophila*-associated *Spiroplasma* strains. The size and icosahedral shape of the particles we recovered from strain NSRO are consistent with those previously described from NSRO and other strains, except that those possessed a short tail. Of the published images of virions from *Drosophila*-associated *Spiroplasma*, short-tails appear to be visible only in those of free virions obtained from direct preparations of fly hemolymph [i.e., not involving density gradient isolation [and Fig. 47 insert 42, 43]]. To our knowledge, no images of density-gradient isolated particles from *Drosophila*-associated *Spiroplasma* have been published, but Cohen et al. [56] stated that they observed short-tailed icosahedrons obtained from particles banded in a metrizamide density gradient. A tail was not visible in intracellular NSRO particles infecting WSRO, or those presumably just extruded from WRSO cells [54]. Similarly, a tail was not visible inside or outside lysing NSRO cells infected with particles from strain *s*Hyd1 [52]. Banding in cesium gradients of similar viruses, such as the one described from *S. citri*, disrupts some particles, resulting in the release of intact tail structures [42]. It is thus possible that our isolation procedure led to the loss of tail structures; a similar phenomenon was hypothesized by Bordenstein et al. [102] to explain the lack of tails in their *Wolbachia* phage preparations.

Several features about the three phage-like contigs we recovered from the particle extractions of NSRO and MSRO-Br (i.e., NSRO-P1, NSRO-P2, and MSRO-P1) are similar to those previously described in *Drosophila*-associated *Spiroplasma* strains. Our “Final” assembly contig sizes of ∼19kbp are within the range of genome sizes from prior reports (17, 21.8, and >30 kpb). The respective sizes of our preliminary NSRO-P1 and NSRO-P2 contigs (31 and 21 kbp), which contained terminal repeats of 12 and 1.3 kbp, were more similar to the > 30 and 21.8 kbp sizes previously reported [56]. Our failure to detect bands consistent with sizes > 5 kbp in the gel is puzzling. It is possible that ∼19 kbp fragments were present, but at concentrations lower than detectable in the gel (the Nanopore library preparation was aimed at enriching fragments > 3 kbp). It is also possible that the conformation of the DNA run on the gels was different from double-stranded and linear, such as supercoiled DNA, which tends to run faster in the agarose gel matrix [103, 104]. Nonetheless, the Nanopore library prep should not have used any ssDNA template because it requires dsDNA to attach the adapters. Prior restriction fragment electrophoresis patterns of phage particles from *Drosophila*-associated strains (only performed on DNA from NSRO and *s*Hyd1 particles) and from non-*Drosophila*-associated strains (i.e., the 21 kpb phage genomes of *S. citri* and *S. mirum*), are consistent with circular permutation and terminal redundancy [44, 56, 105], and thus, expected to possess a headful packaging replication mechanism [106]. The mapping pattern of long and short reads to our phage-like contigs (i.e., artificial concatemer) is also consistent with circular permutation.

Our finding of two distinct phage-like contigs in NSRO (NSRO-P1 and NSRO-P2) is consistent with Cohen et al. [56]’s report of two distinct 21.8 kbp phage genomes based on Restriction Fragment Length Variation. The three phage-like contigs we obtained contain several ORFs with annotation terms associated with phage structural proteins (terminase, portal, head-tail connector, and capsid) and with DNA binding/recombination proteins. Of the latter, the large phage-like terminase annotated in the three phage-like contigs could aid in the packing of viral dsDNA [107], whereas the ssDNA binding protein (in NSRO-P1) and recombinase (in NSRO-P2 and MSRO-P1) would most likely be involved in phage DNA replication [108]. The presence of both types of temperate recombinases, as well as the high (albeit imperfect) homology detected upon mapping NSRO-P1, NSRO-P2, and MSRO-P1 to the main chromosome of the three MSRO-Ug substrains with available draft genomes, suggests that they could be lysogenic. The absence of ORFs encoding recognizable proteins expected in functional phage (e.g. tail structural and lysis proteins), however, raises doubts as to whether they represent genomes of fully functional phages. Nonetheless, it is possible that the necessary genes are encoded among the numerous remaining hypothetical CDS.

Three scenarios illustrated in Figure 5 can explain the recovery of the phage-like and the non-phage-like contigs in our extractions aimed at particles. In Scenario 1, the contigs are packaged into virions, as in the case of fully functional phage or Gene Transfer Agents (GTA). GTAs are particularly common in α-proteobacteria, where random ∼5–15 kbp fragments of chromosomal DNA are packaged into phage-like particles via headful packaging and released via lysis [109-113]. Packaging of non-random regions of the chromosome into GTAs has been reported in *Bartonella bacilliformis*, where run-off replication appears to preferentially amplify a particular region of the genome, which is then packaged into particles [114]. If the contigs we recovered are not packaged into virions, they may represent non-packaged extra-chromosomal (Scenario 2) or non-packaged chromosomal DNA fragments (Scenario 3). A DNAse treatment of particles prior to DNA extraction should help to distinguish scenarios 1 vs. 2 and 3.

**Fig. 5.**
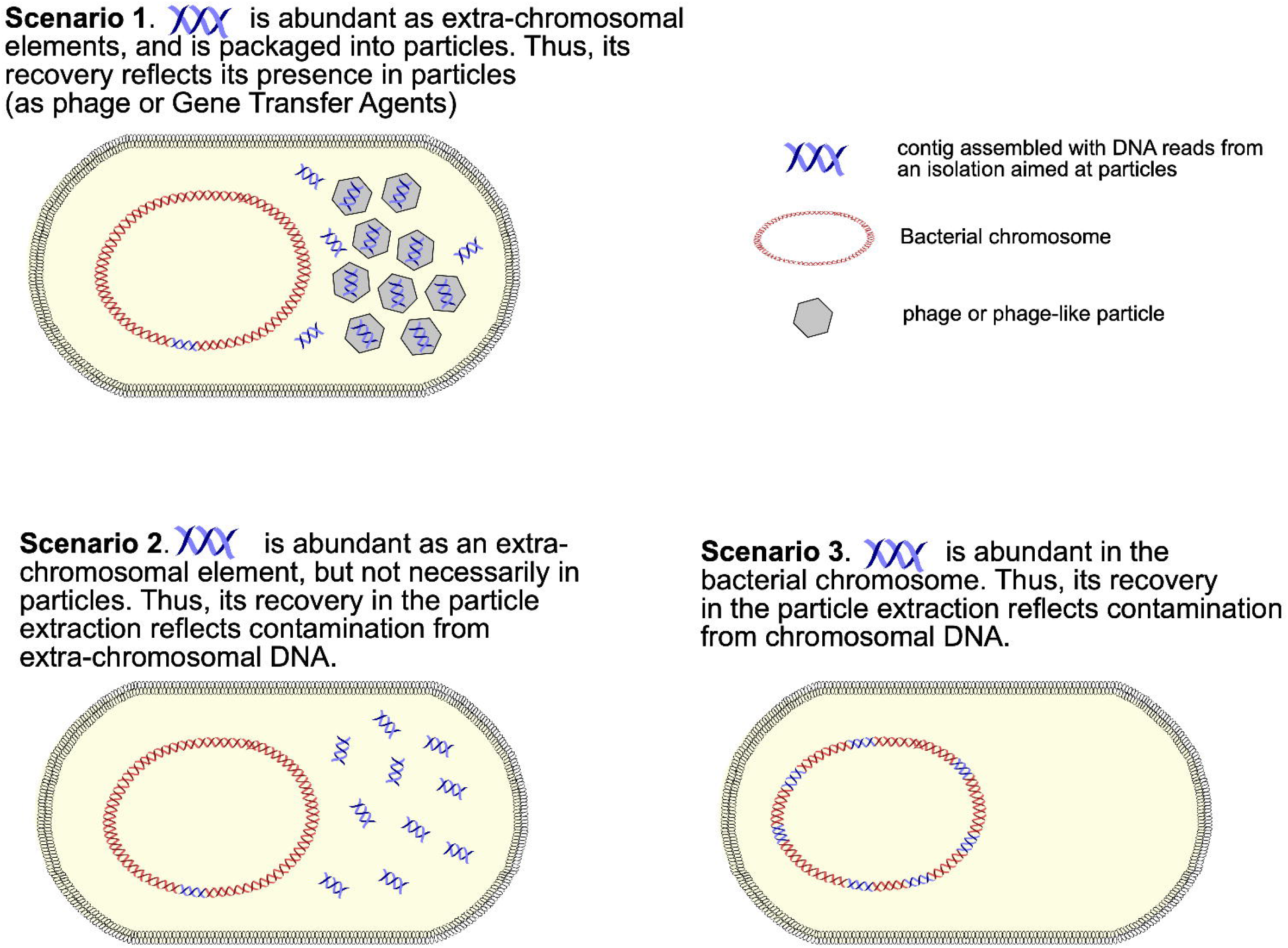
Possible scenarios to explain overrepresentation of a particular locus in a Whole Genome Shotgun sequencing dataset. Cartoon depicting three scenarios that could result in an overrepresentation of a particular DNA sequence (the blue double strand) in an assembly of reads obtained from an isolation aimed at viral or viral-like particles from a sample containing cells of *Spiroplasma*.

Our recovery of the RIP2 gene in the non-phage-like Contig 3 raises intriguing questions given its: (a) ribosome depurination activity; (b) possible implications in killing of parasitic wasps (and nematodes), and *Drosophila* embryos or old adults; (c) high expression levels, and differential expression between conditions in MSRO strains; and (d) differing copy numbers in DNA extracts not enriched for particles (single-copy in MSRO-Ug vs. multi-copy in MSRO-Br; Table 5; unknown in NSRO) [27, 32, 36, 38, 39, 66]. Could having RIP2 DNA copies in virions (Scenario 1) enable horizontal gene transfer? Could having unpackaged extra-chromosomal copies (Scenario 2) or multiple chromosomal copies (Scenario 3) enable higher RIP2 protein synthesis? Could the different RIP2 gene copy numbers detected in published datasets (not from particle-enriched extractions) of MSRO-Br vs. MSRO-Ug be related to the higher degree of overall protection against certain wasps reported for MSRO-Br [115]?

Except for the homology of NSRO-P1, NSRO-P2, MSRO-P1 to UA44405, a region that is deleted in an insect-transmissible strain of *S. citri* [92], and that contains one or more genes annotated as putative adhesins [98], two of which HHPred annotates as possible phage structural genes, we found no evidence that any of the genes encoded could represent “virulence” genes. This contrasts with the findings in *Wolbachia*, where the genes responsible for reproductive manipulation (Cytoplasmic Incompatibility and male killing) reside on WO prophage regions [reviewed in 116]. It is possible, however, that some of the ORFs in the phage-like contigs encode genes that may contribute to the interaction of *Spiroplasma* and its eukaryotic hosts.

The recovery of NSRO-IS5 and MSRO-Br-IS5 contigs in NSRO and MSRO-Br phage particle isolations is also interesting. Both contigs showed high homology to the IS5 family of Insertion Sequences [117]. Insertion Sequence elements are regarded as the shortest mobile elements capable of autonomous transmission, and generally contain an ORF coding for a transposase that is required for transposition [118]. Given the numerous copies of IS5 contigs found in the genome assemblies of *S. poulsonii*, it appears that transposition of this element is frequent and likely contributes to the rapid evolution seen in *S. poulsonii* genomes [27].

## Conclusions and Outlook

Several lines of evidence suggest that we recovered phage-like particles of similar characteristics (shape, size, DNA content) to those previously reported in *Drosophila*-associated *Spiroplasma* strains. Our study is the first one to report sequencing and assembly of DNA isolated from such particles in *Drosophila*-associated *Spiroplasma*. Although the larger (i.e., ∼19 kbp) contigs we recovered contain regions encoding phage-like structural and DNA replication functions, the absence of certain recognizable “essential” phage genes, suggests that they are not fully functional phage. It is worth noting, however, that these contigs encode numerous ORFs annotated as hypothetical, which may encode the “missing essential” phage genes, or unknown “virulence” genes. Even though an efficient system to genetically modify *S. poulsonii* has not been developed [119], the role of *Spiroplasma*-or phage-encoded genes in the interaction with *Drosophila* can be tested by transgene expression in *Drosophila* [e.g. 29, 32]. Our unexpected recovery of smaller contigs, one of which encodes a known and highly expressed toxin gene (RIP2), raises the question as to whether the presence of multiple copies of this gene in the *Spiroplasma* cell (as chromosomal or extra-chromosomal DNA) may contribute to the toxicity of *Spiroplasma* against eukaryotes, and thus influence the *Drosophila-Spiroplasma* interaction.

An important limitation of our study was the low quantity of particles that we were able to isolate, the lack of a chromosomal genome assembly for the specific *Spiroplasma* strain producing the particles, and our inability to cultivate *Spiroplasma* outside its host. Future work would benefit from isolation of substantially larger quantities of particles that could be subjected to diverse assays to determine their protein and DNA content, replication and packaging mechanism, life cycle, and host range.

## Acknowledgments

Anika Stankov for technical assistance. Paul de Figueiredo provided feedback on the research and early versions of the manuscript. Portions of this research were conducted with high-performance research computing resources provided by Texas A&M University (**https://hprc.tamu.edu**). This research was conducted in partial fulfillment of the Master’s of Science degree requirements of Paulino Ramirez.

## Supporting Figure Legends

**Fig. S1 Original Canu assembly of NSRO phage like contigs**. Grey arrows in the reference are imperfect terminal repeats in the contig. A) NSRO-P1, B) NSRO-P2.

**Fig. S2 Alignment of Nanopore reads to the original Canu assembly contigs**. Grey arrows in the reference are imperfect terminal repeats in the contig. The raw long reads are shown in black. Gray outline boxes in reads represent trimmed regions (i.e., different from reference). A) NSRO-P1, B) NSRO-P2.

**Fig. S3 Geneious-based alignments with annotated ORFs of (A) NSRO-P1, NSRO-P2 and MSRO-P1 phage-like contigs, and (B) Insertion element identified in the phage particle assembly from NSRO and MSRO-Br. Sites with identical bases in all positions are depicted green in the identity bar**. In (B) two ORFs were identified along with inverted repeat sequences (grey). The two sequences have 100% homology, except at terminal ends of the contigs. The yellow ORFs (possible translational frameshift) code for a DDE superfamily endonuclease and mobile element like protein respectively, which can form an Insertion element.

**Fig. S4 Alignment of the nucleotide sequences of NCBI U44405.1 and our particle-derived contigs: NSRO-P1, MSRO-Br-P1, NSRO-P2**. Grey = position identical to consensus sequence; black = position different from consensus. Open reading frames (ORFs) are indicated with colored block arrows. The color-coding and labels of our particle-derived contigs follow Fig. 4 (yellow = hypothetical; others assigned a putative function). Yellow annotations in U44405 reflect those in NCBI, whereas the green ones were added as a result of the new findings based on HHPred.

## Supporting Dataset Descriptions

**Supporting Dataset S1**. Four contigs from the MSRO-Br particle extraction that had blast hits to *Spiroplasma*, but whose coverage was extremely low.

**Supporting Dataset S2**. Annotations of the particle-derived phage-like contigs (NSRO-P1, NSRO-P2, Contig3, IS5, and MSRO-P1). Each contig is in a separate sheet. ORFs were predicted and annotated using RastK software. ORFs were then manually annotated with HHPred to elucidate potential function of ORF products. HHPred predictions are described in notes with probability, accession ID and name.

**Supporting Dataset S3. Contamination Report based on Blast Hits**. Top blast hit for each of the contigs recovered from the three sequencing runs: Nanopore and Illumina MiSeq Index 4 for NSRO particle derived DNA (sheets Blast_Index_4_NSRO and Index_4_NSRO_unique_hits); and Illumina MiSeq Index 1 for MSRO particle DNA (sheets Blast_Index_1_MSRO-Br and Index_1_MSRO-Br_unique_hits). The first worksheet contains metadata for each run consisting of information related to potential sources of contamination. Blast output of SPAdes assembled contigs and unique blast hits to organisms are included.

**Supporting Dataset S4. Coverage of the NSROP-1, NSROP-2, Contig3, IS5 and MSROP-1 contigs by published RNAseq reads**. RNAseq read coverage, raw and RPKM, for NSRO and MSRO ORFs are shown in separate sheets.

**Supporting Dataset S5. Repeat detection at the ORF level of particle-derived contings**. Results of RepeatMasker at the Open Reading Frame (ORF) level, for the particle-derived contigs (NSROP-1, NSROP-2, Contig3, IS5 and MSROP-1) against the MSRO-Ug (GCA_000820525.2) and MSRO-Br (GCA_019295385.1) genome assemblies. The first sheet (“Totals”) shows the number of times that each ORF (peg) of the particle-derived contigs was found by RepeatMasker in the genome assemblies. Raw output for each analysis is in a separate sheet.

**Supporting Dataset S6. Repeat detection at the contig level of the particle-derived contigs**. Results of RepeatMasker for the particle-derived contigs (NSROP-1, NSROP-2, Contig3, IS5 and MSROP-1) against ten *Spiroplasma* genome assemblies listed in Table 4.

The first sheet “Legend” describes the header of each column. Raw output for each contig is in a separate sheet.

## Notes

**Conflict of interest** The authors declare that they have no conflicts of interest.

### Competing Interest Statement

The authors have declared no competing interest.

### Summary of Updates

The manuscript has been updated to address the comments from peer-reviewers during its consideration for publication in Current Microbiology

