## Supplementary figures and images for "Preliminary Characterization of Phage-like Particles from the Male-Killing Mollicute *Spiroplasma poulsonii* (an Endosymbiont of *Drosophila*)"

### Figure S1

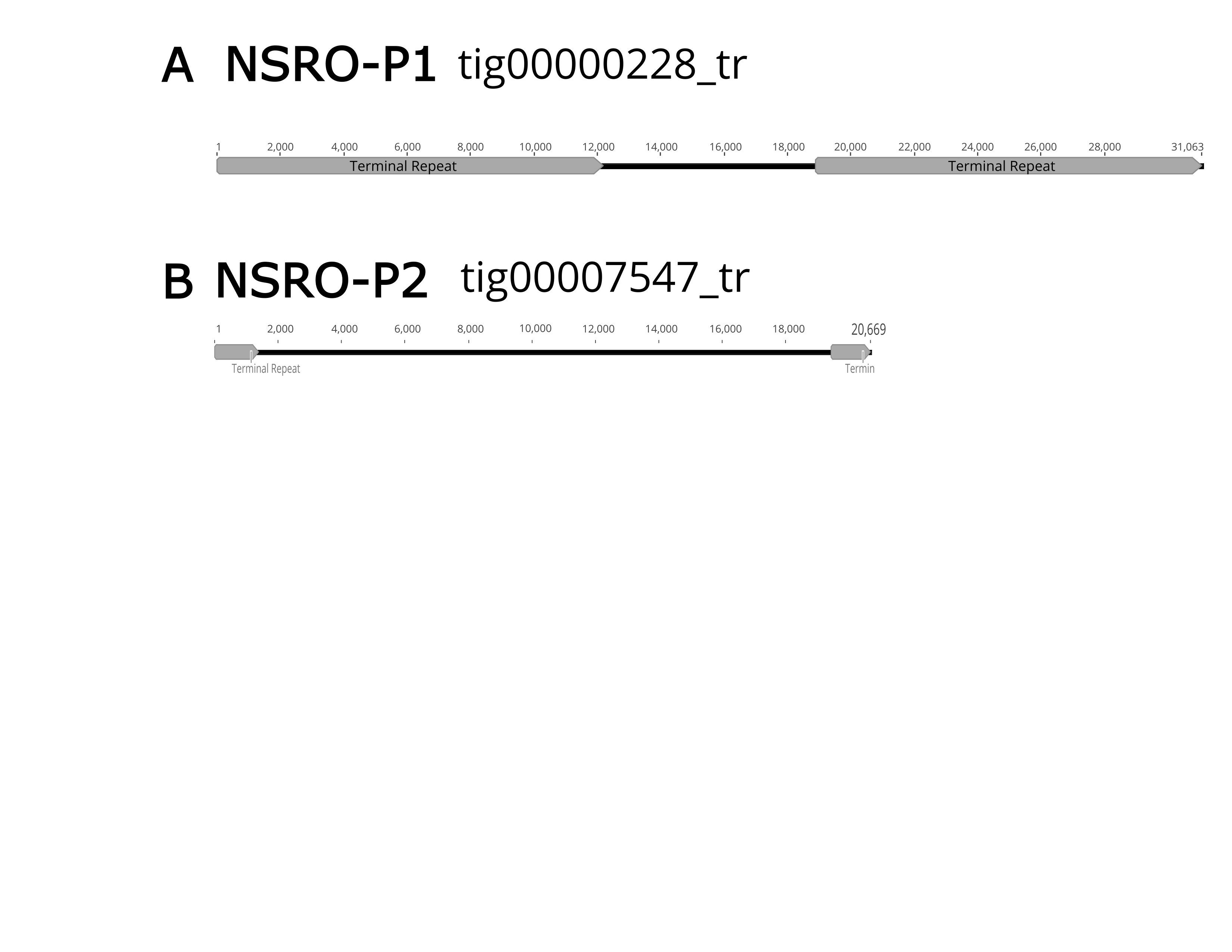

### Figure S2

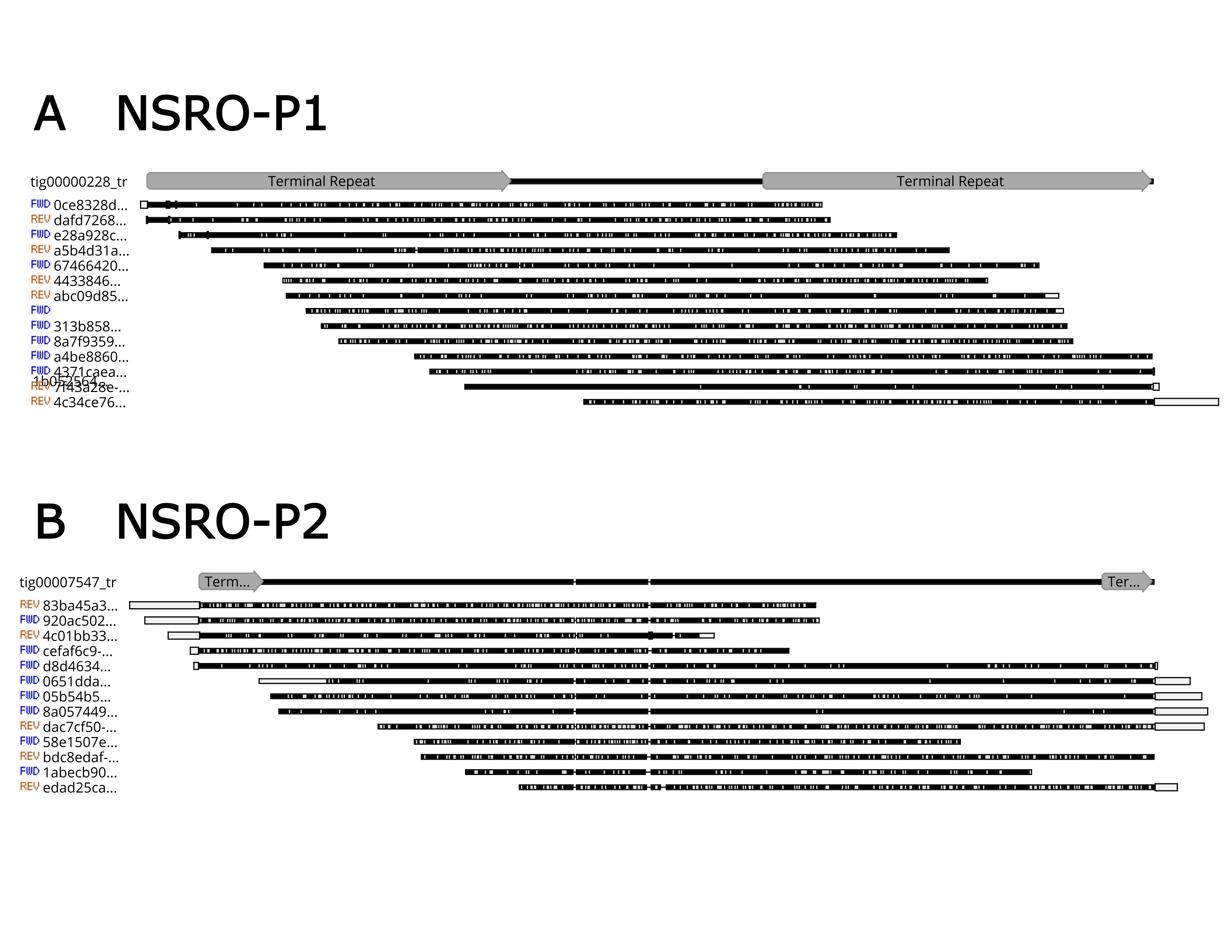

### Figure S3

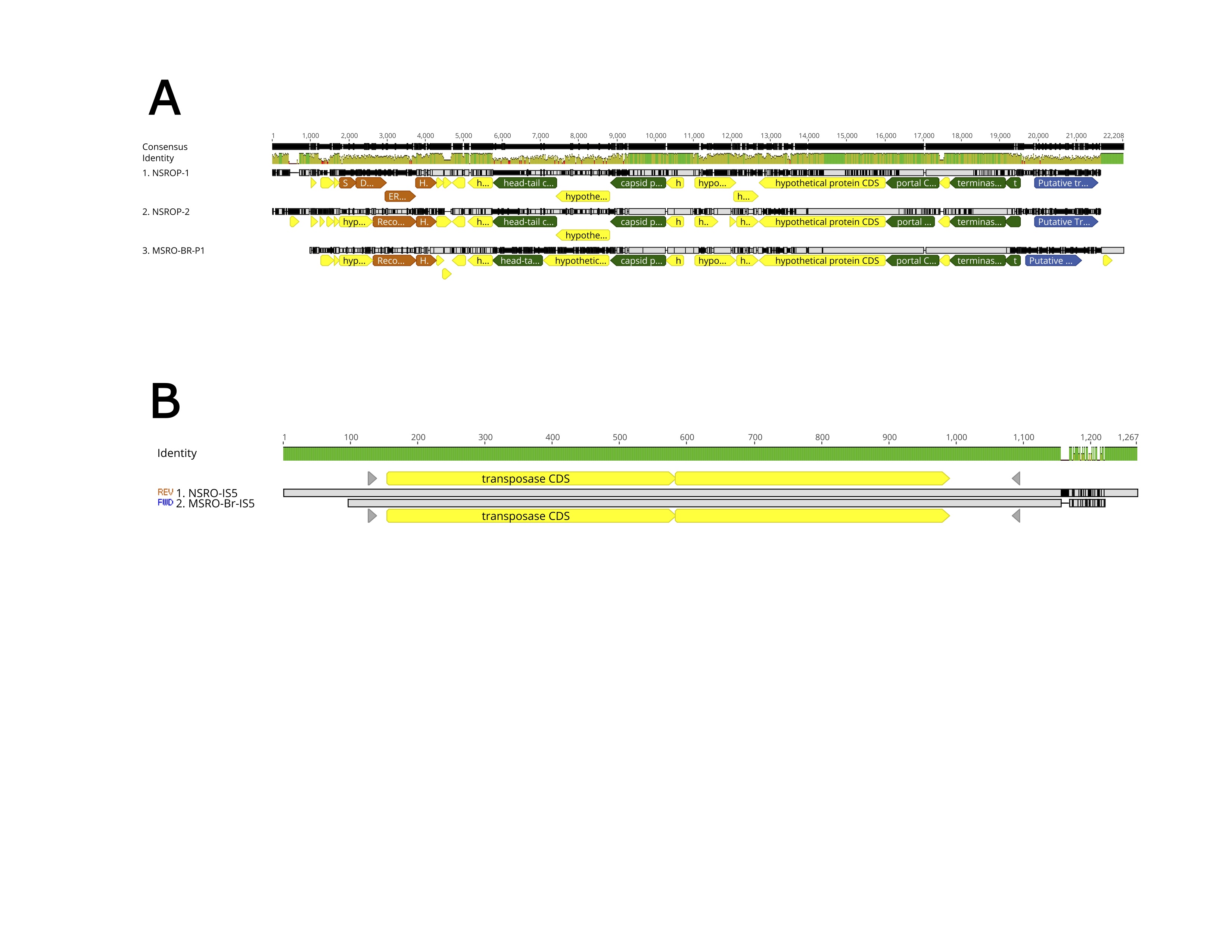

### Figure S4

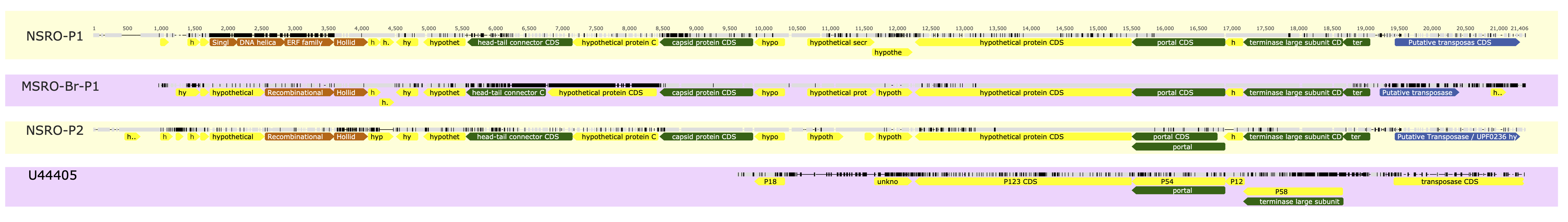
